# Dual roles of idling moments in past and future memories

**DOI:** 10.1101/2024.06.27.600961

**Authors:** Khaled Ghandour, Tatsuya Haga, Noriaki Ohkawa, Chi Chung Alan Fung, Masanori Nomoto, Mostafa R. Fayed, Hirotaka Asai, Masaaki Sato, Tomoki Fukai, Kaoru Inokuchi

**Author notes:** These authors contributed equally to this work (Khaled Ghandour, Tatsuya Haga, Noriaki Ohkawa). Center for Information and Neural Networks (CiNet), NICT (National Institute of Information and Communications Technology), Osaka, Japan. Graduate School of Medicine, The University of Tokyo, Tokyo, Japan (Hirotaka Asai).

## Abstract

Every day, we experience new daily episodes and store new memories. Although memories are stored in corresponding engram cells, how different sets of engram cells are selected for current and next episodes, and how they create their memories, remains unclear. We report that in mice, hippocampal CA1 neurons show an organized synchronous activity in prelearning home cage sleep that correlates with the learning ensembles only in engram cells, termed preconfigured ensembles. Moreover, after learning, a subset of nonengram cells develops population activity, which is constructed during postlearning offline periods through synaptic depression and scaling, and then emerges to represent engram cells for new learning. Together, our findings indicate that during offline periods there are two parallel processes occurring: conserving of past memories through reactivation, and preparation for upcoming ones through offline synaptic plasticity mechanisms.

## Introduction

Our daily experiences form a myriad of memories that accrue to construct our personality. To adequately encode and retrieve these memories, the brain must always be prepared and organized for near-upcoming events and perform parallel processing of past memory. The process of determining which set of neurons is to be recruited for an upcoming event or task is known as memory allocation^1^. Proper allocation of memory governs adequate retrieval of that memory whenever it is needed. Groups of neurons that represent a specific memory are called engram cells^2^. Engram cells can be labeled and identified through the expression of immediate early genes, such as *c-fos*^3,4^. Optogenetical activation or inhibition of these subsets of neurons leads to memory retrieval or silencing, respectively^5-7^. It has been demonstrated that engram cells possess characteristic activity during learning^8,9^. As previous memories affect the next event, previously formed engram cells may play some role in the selection of next engram cells^10-12^. Several previous studies have shown that CREB and neuronal excitability could bias the competition towards the excitable neurons to be an engram^13-16^. However, how these engram cells behave in periods prior to learning and how they affect future learning, is yet to be studied. On the other hand, hippocampal intrinsic dynamics form a repertoire of preexisting assemblies of high temporal sequential events ahead of time^17-19^ forming a network of preconfiguration, suggesting that coding principles in neuronal networks might reflect selection of remarkably excitable neurons from much larger groups of properly connected cells.

Sleep’s contribution to memory consolidation has been extensively investigated in recent decades. Brain activities occurring during offline periods when subjects are not engaged in a certain task, such as during sleep or awake rest, are defined as “idling state” activities^20,21^. Neuronal reactivation and replay are responsible for the consolidation and strengthening of acquired memories^8,22,23^. Artificial inhibition or disruption of theta waves or sharp wave ripples (SWRs) during either rapid-eye-movement (REM)^24^ or non-REM (NREM)^25^ sleep, respectively, disrupts the retrieval of recent memories. However, sleep is now thought of as an active process that is required for more than just consolidation of past memories, rather than a passive process ^23^. Sleep and awake rest reactivations are linked with many behavioral outcomes like learning^26^, creativity^27-29^, and planning, either toward a reward^30^ or away from an expected averse outcome^31^. Although the concept of sleep has recently expanded, with sleep now being considered a more dynamic process of diverse functions, the physiological role of sleep in memory acquisition and encoding is not well understood.

Episodic memories do not merely reflect past experiences but are also important for thinking about the future^2^. Several functional magnetic resonance imaging (fMRI) studies provide evidence^32,33^ that denotable activity associated with the past occurs in the brain when thinking about the future, mimicking mental time travel^34^. This high overlap between past and future events is in agreement with findings where hippocampal lesions or damages lead to an inability in thinking and imagining about the past and future, respectively^35,36^. Nonetheless, how and when our brains prepare and modify our existing memories to accommodate our new experiences remains unclear. In this study, we addressed the questions of whether and how sleep plays a role in the selection of engram cells.

## Results

### Active ensembles during prelearning sleep are recruited into upcoming engram

To understand how a specific set of neurons are chosen to represent a particular event as a memory trace, we need to track the activity of engram cells across different memory processing stages. We recently established a Ca^2+^ imaging system using a miniature microscope^8^ that is suitable for visualizing both engram and nonengram cells across memory processing stages in freely moving mice^37,38^ (Fig. 1A, Supp Fig. 1). We used double transgenic Thy1-G-CaMP7 mice crossed with c-fos-tTA mice and the miniature microscope to track and label the activity of both engram and nonengram cells. Lentivirus (LV), harboring KikGR (coding for Kikume Green Red fluorescent protein)^39^ under the control of the tetracycline responsive element (TRE), was injected into the hippocampal CA1 region to label the engram CA1 cells, with the result that activated cells express KikGR protein in the absence of doxycycline (DOX; Fig. 1A). We previously showed that cells labeled by this system represent engram cells because artificial activation of these cells triggers a memory recall^8^. KikGR is irreversibly photoconverted from green to red fluorescent protein by exposure to UV light of 365 nm (Fig. 1B), which allows simultaneous Ca^2+^ imaging with G-CaMP7. EEG and EMG electrodes were implanted along with the microscope to allow different sleep stages to be differentiated (Suppl. Fig.2A and see Methods). Imaging on day 1 was performed in prelearning NREM (N1) and REM (R1) sleep, during learning (A), and in postlearning NREM (N2) and REM (R2) sleep. This was followed by imaging sessions during retrieval (A’) and in a different context (B) on day 2 (Fig. 1B). Calcium transients were detected automatically using a HOTARU online sorting system^40^. All detected cells were categorized into engram and nonengram cells according to a snapshot image taken 24 hours after the context exposure (Suppl. Fig. 2B-E). Engram cells constituted 7 % (±1.5) of the total cells recorded while nonengram cells constituted 93% (±1.5) (Suppl. Fig. 2F and Suppl. Table 1).

**Figure 1:**
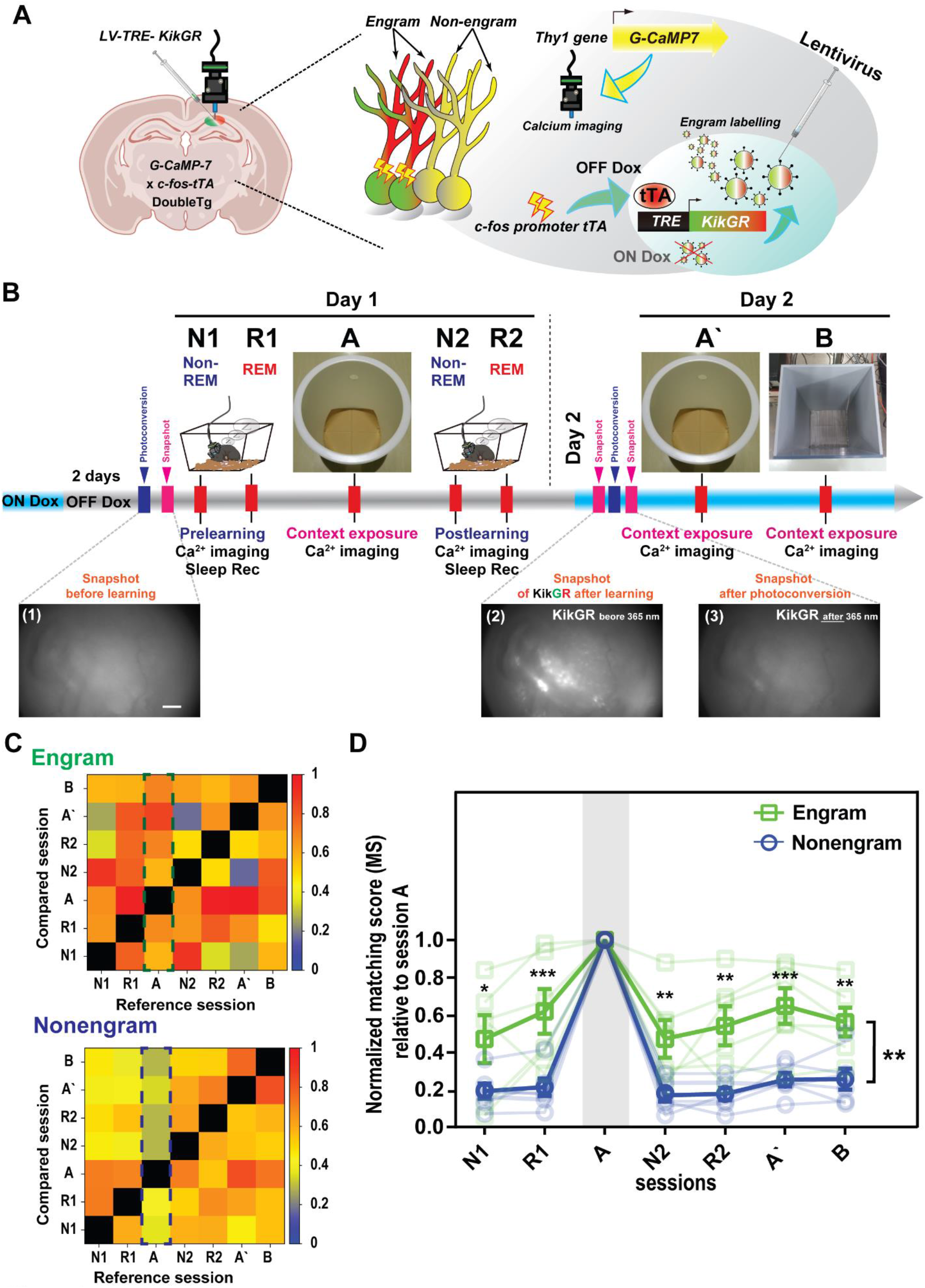
Ensembles activated during prelearning sleep are recruited into the upcoming engram. (A) Schematic diagram showing the miniature microscope placed at the hippocampal CA1 in c-fos-tTA x G-CaMP-7 double transgenic mice injected with TRE-KikGR lentivirus for labeling and visualizing both engram and nonengram cells. (B) Experimental design for the calcium imaging and behavior paradigm. NREM (N1) and REM (R1) in prelearning sleep sessions, and then learning (A), are followed by postlearning sleep sessions NREM (N2) and REM (R2) on day 1. On day 2, a snapshot for KikGR is taken, and then after ∼2 hours, a retrieval session (A’) was followed by a different context (B) (top), Representative snapshots of KikGR (scale bar, 100 µm) taken before learning (1), 24 hours after learning (2), and after photoconversion to render KikGR invisible to the microscope (3) (bottom). (C) Representative matching score (MS) matrixes for engram (top) and nonengram (bottom) cells across sessions to show similarity between ensembles. (D) Normalized MS analysis in reference to session A for both engram (green) and nonengram (blue) cells across sessions. n = 6, statistical comparisons were made between engram and nonengram using two-way repeated-measures ANOVA, F (1, 5) = 36.73, p = 0.0018 (engram vs nonengram), Bonferroni’s multiple comparisons test, p = 0.0187 (N1), p = 0.0003 (R1), p = 0.0091 (N2), p = <0.0013 (R2), p = <0.0004 (A’), p = <0.008 (B). ^*^p < 0.05, ^**^p < 0.01, ^***^p < 0.001. Data represent the mean ± s.e.m.

**Figure 2:**
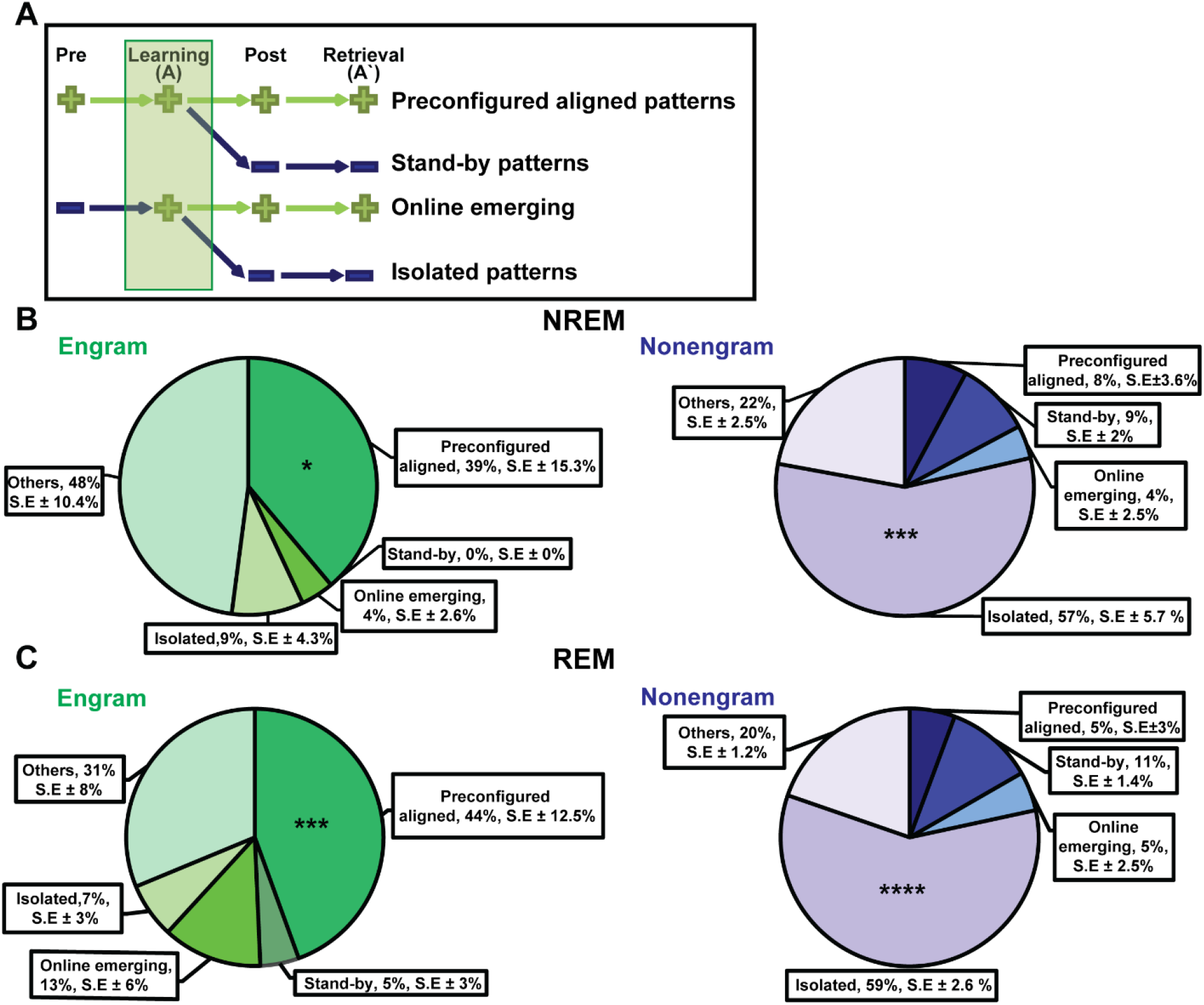
Preconfigured ensembles are aligned across both pre- and postlearning sleep in engram cells. (A) Schematic diagram of the tracking of all learning ensembles with ensembles detected in other sessions (pre-, postlearning sleep, and retrieval). Ensemble pairs have positive marks (+) or negative marks (−) indicating activation or absence of that ensemble, respectively. Ensembles were considered active if the ensemble pair, from a different session, had normalized cosine similarity >0.6. (B and C) Pie charts showing percentage of ensembles relative to the learning session (A) using the NREM sleep only (B) or the REM sleep only (C) in pre- and postlearning sleep. (B) n = 6, statistical comparisons were made between engram and nonengram using two-way repeated-measures ANOVA, F (1, 5) = 0.2415, p = 0.643 (engram vs nonengram), Sidak’s multiple comparisons test, p = 0.0180 (preconfigured aligned), p = 0.8649 (stand-by), p > 0.99 (online emerging), p = < 0.0003 (isolated), p = 0.0622 (others). (C) n = 6, two-way repeated-measures ANOVA, F (1, 5) = 5.841, p = 0.0603 (engram vs nonengram), Sidak’s multiple comparisons test, p = 0.0002 (preconfigured aligned), p = 0.9308 (stand-by), p = 0.8496 (online emerging), p = < 0.0001 (isolated), p = 0.508 (others). ^*^p < 0.05, ^***^p < 0.001, ^****^p < 0.0001. Data represent the mean ± s.e.m.

Correlation matrix repetition analysis showed that engram cells possess characteristically higher repetitive correlation during contextual learning (Suppl. Fig. 2G)^8^. We previously described a method that is well suited for picking neurons that are activated together and their activity patterns^8^, non-negative matrix factorization (NMF) analysis^41,42^. In brief, the population activity ensembles were selected on the basis of the Akaike information criterion with a second-order correction (AICc), in which the data matrix was approximated by two non-negative factors (the pattern matrix and the intensity of that pattern activity) to calculate an error function (Suppl. Fig. 3A). Collectively, NMF analysis decomposed the total complex pattern activity into a series of group activated neurons (termed ensembles) and their temporal activity patterns (Suppl. Fig. 3B and see Methods). Each ensemble is constituted of different individual cells’ contributions and their grouped synchronous activity across different time points; therefore, several ensembles have different cell compositions and temporal activity. To track the activity of the detected ensembles across other sessions, we applied the normalized dot product (cosine similarity) to compare the similarity between all pairs of pattern vectors of ensembles across two different sessions. If two ensemble vectors completely share the cell identities and activity, the normalized dot product score is 1. However, if there is no similarity, the score is 0. In this study, an ensemble was considered active in another session if it had a normalized dot product equal to or above 0.68, below 0.6 was considered a weak threshold (Supp. Fig. 3C). Comparable results were obtained regardless of higher thresholds. NMF analysis was applied to all recorded sessions (session N1 to session B) for all cells, and separately to engrams and nonengrams (Suppl. Fig. 3D-G). Then, to estimate the reoccurrence of the ensembles across the sessions, we calculated the matching score (MS), which shows the fraction of ensembles from session X that is similar to any of the ensembles from session Y (see Methods; Fig. 1C). The normalized MS results from session (A) were quantified and compared with those of all other sessions in both engram and nonengram cells. Both engram and nonengram ensembles were normalized to their shuffled engram and nonengram ensembles, respectively. Collectively, similar neuronal ensembles were repeated more often across different sessions in engram cells. By contrast, very few ensembles detected in nonengram cells were repeated (Fig. 1D). Engram cells showed significantly higher normalized MS than nonengram cells across all sessions, exhibiting around 40–60% resemblance, even before the learning in prelearning sleep sessions (N1 and R1). Given the fact that engram cells constitute only a small proportion of the total recorded cells, it is important to make sure that the difference found in MS activity was not attributed to the difference in cell size number. We re-performed the dot product and MS analysis using the same number of cells for both nonengram and engram. Engram cells still show higher MS values, and the same conclusion is maintained (Suppl. Fig. 4A and see Methods). This is consistent with our previous finding8 that engram cells with high MS results maintained their high activity in replay (N2 and R2) retrieval (A’) sessions, and that the engram ensembles decreased significantly from the retrieval session (A’) to a different context (B; Suppl. Fig. 4B).

**Figure 3:**
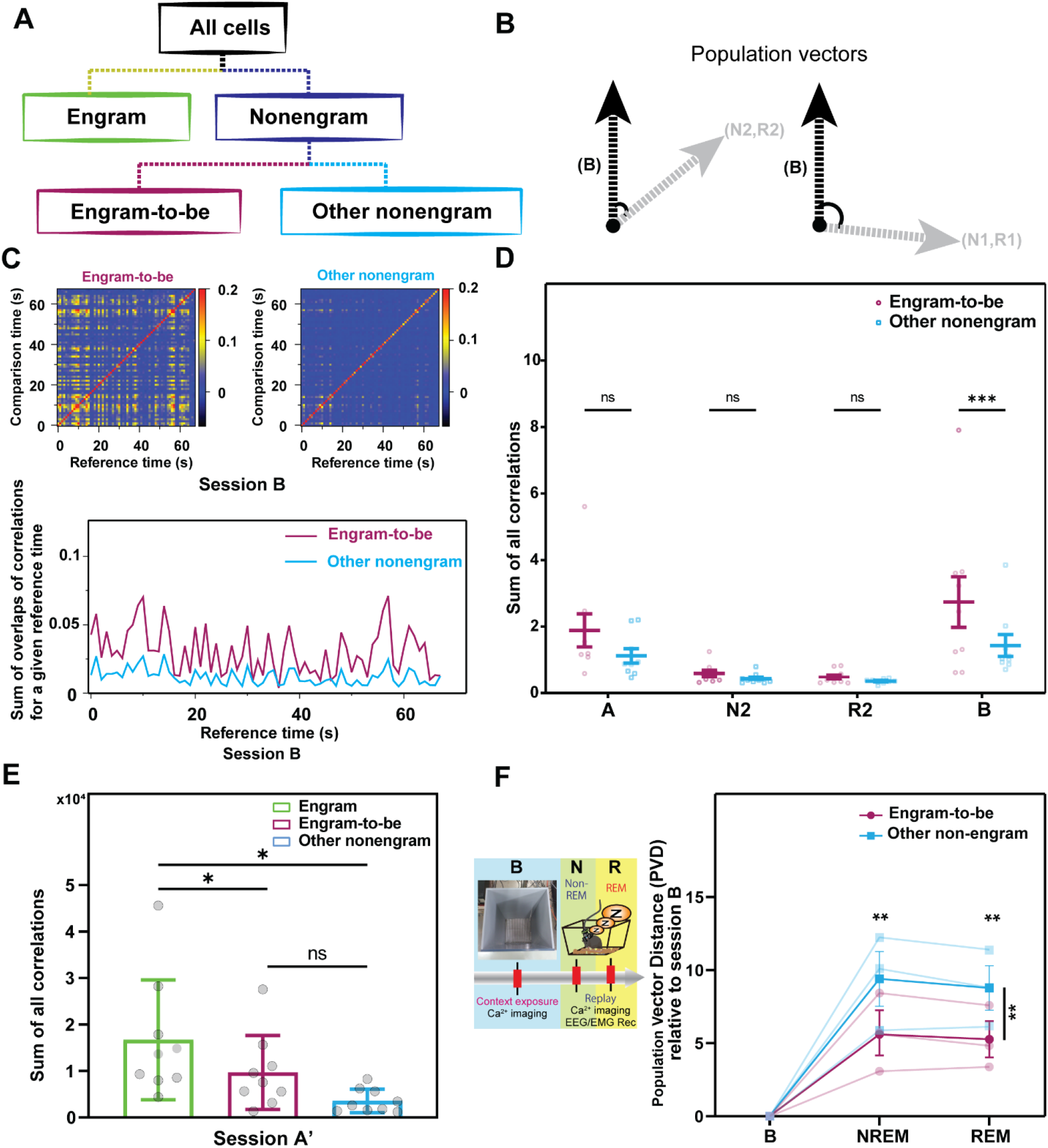
Engram-to-be cells, chosen from the nonengram pool, act as future engram cells for upcoming events. (A) Phylogenetic tree diagram showing classification of all cells into engram and nonengram cells. Nonengram cells are then further subdivided into engram-to-be and other nonengram cells. (B) Illustration showing the first basis for choosing engram-to-be cells. Cells constituting the ensemble of session B having high normalized cosine similarity (>0.6) with the ensemble from sessions N2&R2 (left) and a low normalized cosine similarity (<0.6) with sessions N1&R1 (right). (C) Correlation matrix overlaps between two time points M (t, t’) for the first 60 seconds of session B (See Methods) for engram-to-be (left) and other nonengram (right), and the temporal sum of overlaps at a given time point Mtot(t) (bottom) from a given representative animal (see Methods). (D) Sum of all correlations (Mtot(t)) in the first 60 seconds for engram-to-be and other nonengram across sessions. Statistical comparisons were made using two-way repeated-measures ANOVA, n = 9, F (1, 8) = 5.954, p = 0.0406 (engram-to-be vs other nonengram), Bonferroni’s multiple comparisons test, p = 0.0527 (A), p > 0.99 (N2), p > 0.99 (R2), p = 0.0005 (B). (E) Sum of all correlations for engram-to-be and other nonengram in session A’. Statistical comparisons were made using one-way repeated-measures ANOVA, n = 9, F = 10.53, p < 0.0055 (between engram, engram-to-be, and other nonengram). Tukey’s multiple comparisons test, p = 0.0484 (engram vs engram-to-be), p = 0.0193 (other nonengram vs engram), p = 0.0525 (other nonengram vs engram-to-be). (F) PVD in both NREM and REM with respect to the learning session (B) in engram-to-be and other nonengram cells. Mice were subjected to sessions N1 to B, and then to postlearning B sleep sessions (NREM and REM). n = 3, two-way repeated-measures ANOVA, F (1, 2) = 68.93, p = 0.0142 (engram-to-be vs other nonengram), Sidak’s multiple comparisons test, p = 0.002 (NREM), p = 0.0022 (REM). ^*^p < 0.05, ^****^p < 0.0001, n.s., nonsignificant. Data represent the mean ± s.e.m.

**Figure 4:**
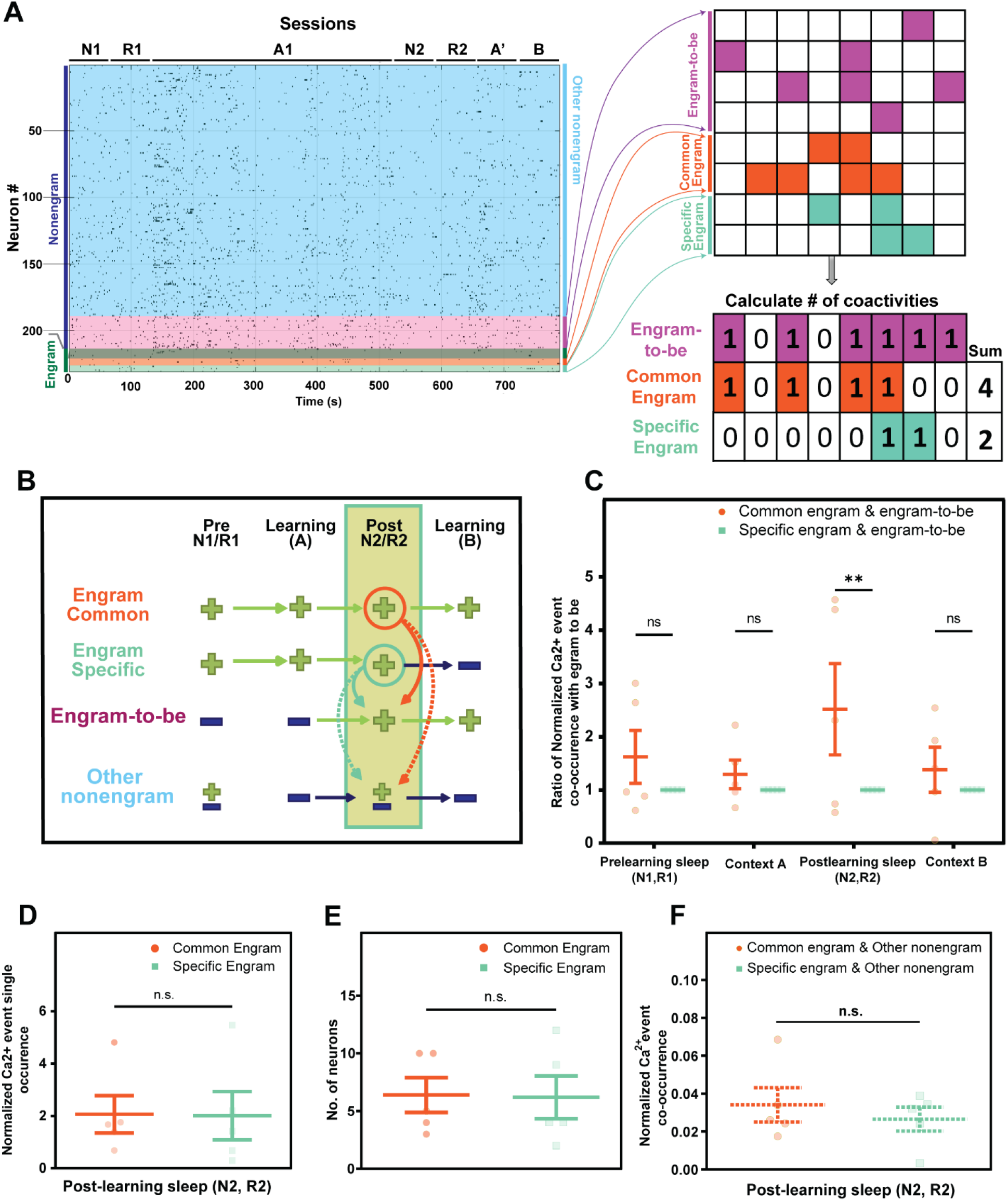
Common engram cells are coactivated with engram-to-be cells during postlearning sleep. (A) Raster plot for all detected neurons of a representative animal across all sessions (left). Magenta, yellow, and light green blocks represent activity (time bin, 250 ms) of engram-to-be, common engram, and specific engram cells, respectively (top right). Schematic diagram of activity in each block and total coactivities between neuron groups (bottom right). (B) Schematic diagram of engram subclassification into common engram and specific engram cells based on their activity in session B, engram-to-be, and other nonengram cells. Similar to Fig. 2A, positive (+) and negative (−) marks indicate ensemble pair similarity and dissimilarity across sessions, respectively. Circles and arrows indicate total coactivities calculated between subtypes. (C) Normalized calcium event co-occurrences between common engram and engram-to-be, compared with specific engram and engram-to-be across all sessions. Statistical comparisons were made using two-way repeated-measures ANOVA, n = 5, F (1, 4) = 2.487, p = 0.1899 (common engram and engram-to-be vs specific engram and engram-to-be), Bonferroni’s multiple comparisons test, p = 0.5230 (prelearning sleep), p > 0.99 (A), p = 0.0076 (postlearning sleep), p > 0.99 (B). (D) Normalized single calcium events of common engrams and specific engrams during postlearning sleep. Statistical analysis was performed using a two-tailed t test between common engrams and specific engrams, n = 5, t = 0.1939, p = 0.8557. (E) Normalized calcium event co-occurrences between common engrams and other nonengrams compared with specific engrams and other nonengrams during postlearning sleep. Statistical analysis was performed using a two-tailed t test between common engrams and specific engrams, n = 5, t = 0.7450, p = 0.4977. (F) Number of neurons of common engrams and specific engrams. Statistical analysis was performed usin g a two-tailed t test between common engrams and specific engrams, n = 5, t = 0.09713, p = 0.9273. ^**^p < 0.01, n.s., nonsignificant. Data represent the mean ± s.e.m.

To check the necessity of sleep in this process, on another cohort of mice, we added awake (Aw) and sleep (S) sessions 1 day before learning (Day 0; Suppl. Fig. 4C). Remarkably, engram cells show highly correlated ensemble activity in prelearning periods only during sleep (S1), but not awake (Aw), sessions when compared to learning session A (Suppl. Fig. 4D). Moreover, engram cells show highly correlated activity even 1 day prior to learning, only during sleep session (S0). This indicates that a group of preconfigured neuronal ensembles that are prominent during sleep, but not awake, sessions are recruited into an upcoming engram upon exposure to contextual learning, and such activity is specific to engram cells.

### Preconfigured ensembles constitute around half of engram ensembles

Then, to know the fate and lifetime of each ensemble detected, we tracked the activity of all ensembles detected in A across sessions (Fig. 2A). All ensembles detected in learning were then named and classified based on their activity in other sessions. Briefly, ensembles activated before and continued their activity after learning were termed ‘preconfigured aligned,’ while ensembles activated before and during learning but were not correlated with any of the ensembles after the learning were termed ‘stand-by’ ensembles. Alternatively, ensembles active during and after learning but were not correlated with any of the ensembles active before the learning were termed ‘online emerging’ ensembles. Finally, ensembles that were not active either before or after learning were termed ‘isolated’ ensembles (Fig. 2A). To simplify the analysis, ensembles that did not match any of the previously mentioned patterns of activity were termed as ‘others.’ We applied this tracking to all detected ensembles of engram and nonengram. The preconfigured-aligned ensembles comprised around 40% of all engram ensembles when NREM sleep (N1 and N2) was used for activation during pre and postlearning sleep (Fig. 2B), whereas isolated ensembles were most abundant in nonengram ensembles (around 60% of ensembles). Similar results were obtained when REM sleep (R1 and R2) was considered (Fig. 2C). This shows that neuronal ensembles that are activated together, even before a certain event, constitute a hippocampal engram and are later activated in postlearning sleep and retrieval sessions. This offline reactivation, especially in engram ensembles, maybe representing what is known as the consolidation process for this event.

### Engram-to-be ensembles arise from nonengram population to represent future memory

Memory allocation theory holds that neurons having relatively higher excitability would be the memory bearers and the winners of the allocation competition^1^. However, it remains unclear what happens to the neurons that do not allocate the memory (losing neurons) in upcoming events, in this case the nonengram cells. Consequently, we investigated whether there is a temporal shift from random to coordinated pattern activities in these nonengram cells across the memory processing stages. We searched the pool of nonengram cells for a group of neurons that could possibly become engram cells in the upcoming event (B) (Fig. 3A).

We sorted all the nonengram cells. First, on the basis of the characteristic features of the engram cells showing preconfigured aligned ensemble activity in the sleep sessions before learning, we searched for ensembles in session (B) that showed high cosine similarity with ensembles in postlearning sleep (N2 and R2) but not prelearning sleep (N1 and R1) sessions (Fig. 3B and Suppl. Fig. 5A). These candidate ensembles were picked up as engram-to-be ensembles (Suppl. Fig. 5B and C). Next, we picked up actively participating cells in these ensembles and termed them “engram-to-be cells”. In principle, engram-to-be cells are cells that are active as an ensemble in session B and show correlated activity in sleep sessions N2 and R2, but not in sleep sessions N1 and R1 before the first learning exposure.

**Figure 5:**
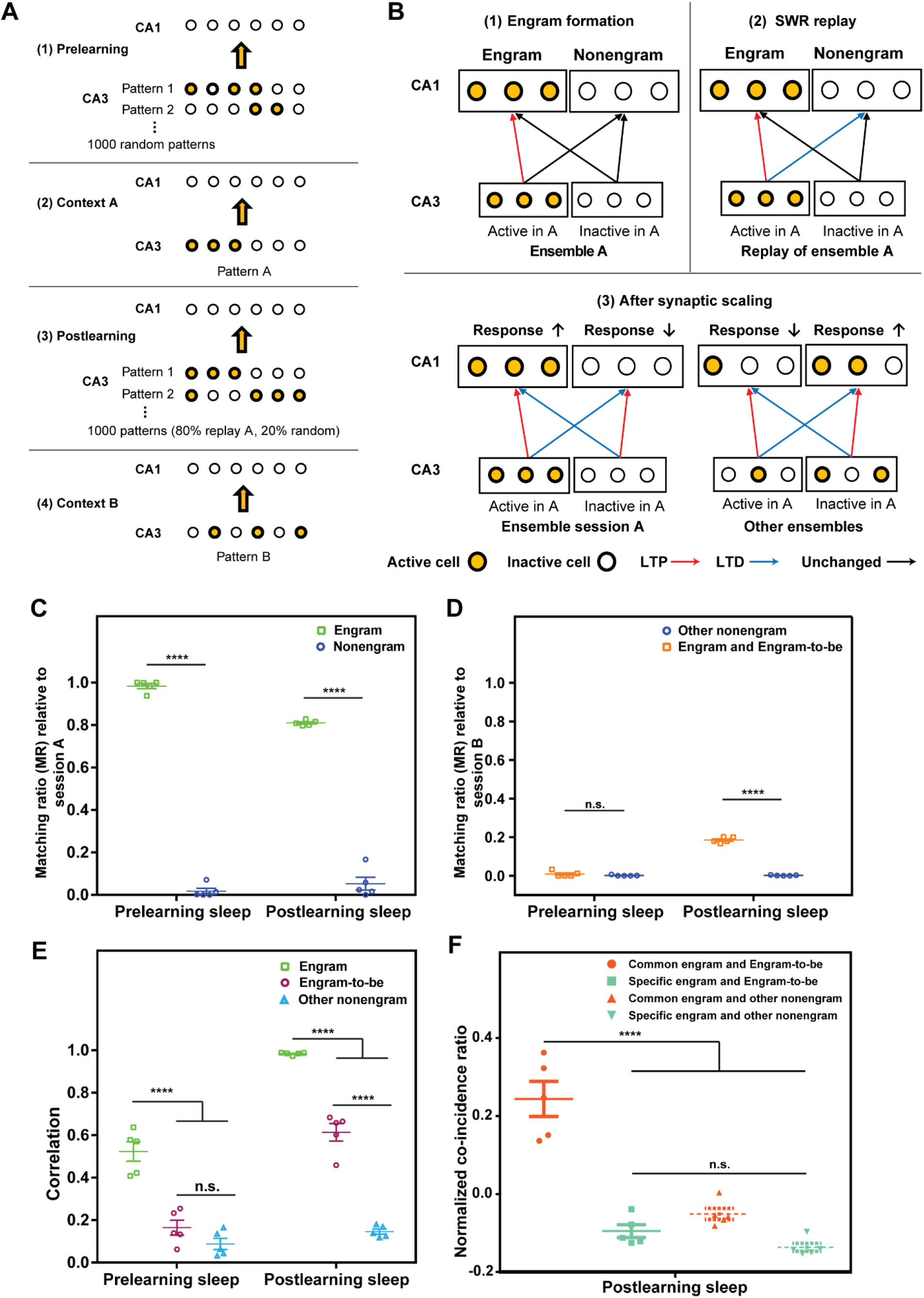
A neural network model reproduces emergence of engram-to-be cells and coactivity with common engram cells. (A) Schematic diagram of the neural network model sessions; (1) prelearning sleep, (2) context A, (3) postlearning sleep, and (4) context B. Yellow and hollow dots symbolize active and inactive cells, respectively. (B) Illustrative diagram of the neural model showing engram formation during learning (1), the plasticity mechanism occurring through sharp-wave ripples during sleep (2), and weight updating by synaptic scaling (3). Red, blue, and black arrows indicate long-term potentiation (LTP), long-term depression (LTD), and no change, respectively. (C) The matching ratio between session A and pre- or postlearning sleep in engram and nonengram cells. n = 5, statistical comparisons were made between engrams and nonengrams using two-way repeated-measures ANOVA, F (1, 4) = 2022, p < 0.0001, Sidak’s multiple comparisons test, p < 0.0001 (prelearning sleep), p < 0.0001 (postlearning sleep). (D) The matching ratio between session B and pre- or postlearning sleep in engram and engram-to-be, and other nonengram cells. n = 5, statistical comparisons were made between engram and engram-to-be vs other nonengram using two-way repeated-measures ANOVA, F (1, 4) = 275.4, p < 0.0001 (cell types), Sidak’s multiple comparisons test, p = 0.1530 (prelearning sleep, engram and engram-to-be vs other nonengram), p < 0.0001 (postlearning sleep, engram and engram-to-be vs other nonengram). (E) Pairwise correlations among engram, engram-to-be, and other nonengram cells in both pre- and postlearning sleep. n = 5, statistical comparisons were made between engram, engram-to-be, and other nonengram using two-way repeated-measures ANOVA, F (2, 8) = 97.32, p < 0.0001 (engram vs nonengram), Tukey’s multiple comparisons test, p < 0.0001 (prelearning sleep, engram vs engram-to-be), p < 0.0001 (prelearning sleep, engram vs other nonengram), p < 0.0626 (prelearning sleep, engram-to-be vs other nonengram), p < 0.0001 (postlearning sleep, engram vs engram-to-be), p < 0.0001 (postlearning sleep, engram vs other nonengram), p < 0.0001 (postlearning sleep, engram-to-be vs other nonengram). (F) The normalized coincidence ratio during postlearning sleep. n = 5, statistical comparisons were made between common engram and engram-to-be, specific engram and engram-to-be, common engram and other nonengram, and specific engram and other nonengram using one-way ANOVA, F (3, 16) = 45.8, p < 0.0001, Tukey’s multiple comparisons test, p < 0.0001 (common engram and engram-to-be vs specific engram and engram-to-be), p < 0.0001 (common engram and engram-to-be vs common engram and other nonengram), p < 0.0001 (common engram and engram-to-be vs specific engram and other nonengram), p = 0.6226 (specific engram and engram-to-be vs common engram and other nonengram), p = 0.6669 (specific engram and engram-to-be vs specific engram and other nonengram), p = 0.1238 (common engram and other nonengram vs specific engram and other nonengram). ^****^p < 0.0001, n.s., nonsignificant. Data represent the mean ± s.e.m.

To ensure that the engram-to-be ensembles represented the majority of the potential engram ensembles of the next event, we applied these criteria to the engram ensembles of session A (Suppl. Fig. 5D and E). The majority of engram ensembles that were active before learning continued to be active in postlearning sleep and retrieval sessions (Fig. 2, Suppl. Fig. 4D). Given that the engram ensemble is defined as those activated during both learning and retrieval, true engram-to-be ensembles (patterns 1 and 3) formed the majority of the predicted engram-to-be ensembles that were identified by the above criteria, whereas pseudopositive engram ensembles (patterns 2 and 4) formed only a small percentage (Suppl. Fig. 5D). Similarly, the true engram-to-be ensembles that were missed by the criteria (false-negative engram-to-be ensembles, patterns 5 and 7) comprised only a small proportion (Suppl. Fig. 5E). This indicates that the criteria applied to pick up the engram-to-be ensembles did indeed capture the majority of meaningful ensembles.

Next, we calculated the correlation repetition analysis for the engram-to-be cells compared with the other nonengram cells, to check if they have the same feature of engram cells during learning. The engram-to-be cells showed higher correlated matrix activity compared to other nonengram cells, denoted by their synchronous repetitive activity and recurring temporal pattern of activity across different time points within a session (session B; Fig. 3C). When the repetitive correlated activity was quantified, only the contextual learning in session B in the engram-to-be cells was significantly higher than in the other nonengram cells (Fig. 3D), while other sessions were comparable. This activity is similar to how the engram cells were behaving in session A (Suppl. Fig. 2G). However, both engram-to-be and other nonengram showed lower correlated activities in session A’ compared to engram cells (Fig. 3E), indicating that the temporal shift and increase in engram-to-be cells activity was specific to new learning (B). The engram-to-be cells showed higher basal activity than the other nonengram cells in session B (Suppl. Fig. 6A). In terms of population size, engram and engram-to-be populations both had comparable numbers of cells, and both were considerably smaller than other nonengram populations (Suppl. Fig. 6B and Suppl. Table 1). On the cellular level, around half (44%) the engram-to-be cells had significantly increased activities in session B compared with session A (Suppl. Fig. 6C and E), while only 37% of the other nonengram population showed a significant increase activity in session B (Suppl. Fig. 6D and E). To assess the necessity of the postlearning sleep (N2 and R2) ensembles activation, we picked up neurons constituting patterns that are not active in N2 and R2 sleep but active during session B and termed them isolated B cells (Suppl. Fig. 7A). Then, we compared the activity of engram-to-be with isolated B cells during session B. Correlated activity of engram-to-be cells were significantly higher during session B than the isolated B ones (Suppl. Fig. 7B), denoting the importance of sleep activation for the engram-to-be preparation and next learning. Overall, these results indicate that there is a subset of nonengram cells that have drastically altered their activity after the 1^st^ learning session to suit the next upcoming learning. Furthermore, we aimed to investigate whether there was a shift in the overall activity of the nonengram cell population through an unsupervised analysis (Suppl. Fig. 8). We conducted a PCA-ICA analysis on the entire nonengram cell population and monitored activities of the ensembles identified during the postlearning sleep session (N2 and R2) in relation to other sessions, prelearning sleep, A, A’, and B. Notably, the ensembles from the postlearning sleep session exhibited a significantly higher level of activities during session B when compared to their activities during the prelearning period. This observation further confirms a notable alteration in the overall population activity patterns during postlearning sleep, patterns that were absent during the prelearning sleep.

**Figure 6:**
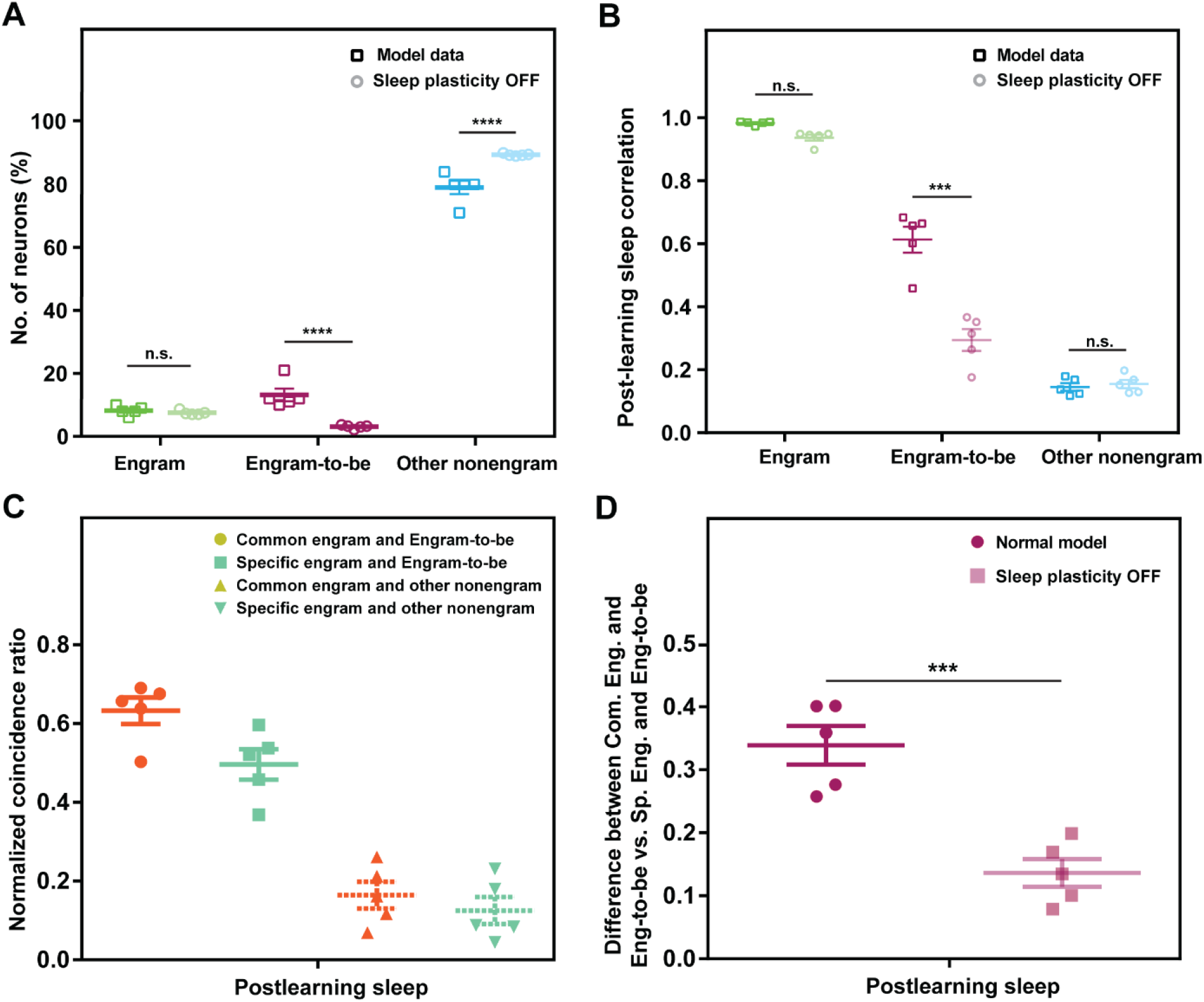
Abolishing sleep plasticity in the network model hinders emergence of engram-to-be cells. (A) Percent of neurons of engram, engram-to-be, and other nonengram cells in the normal plasticity and sleep plasticity OFF models. n = 5, statistical comparisons were made between the normal model and sleep plasticity OFF model using two-way ANOVA, F (2, 24) = 2532, p < 0.0001 (model data vs sleep plasticity OFF), Sidak’s multiple comparisons test, p = 0.9709 (Engram), p < 0.0001 (engram-to-be), p < 0.0001 (other nonengram). (B) Comparing the pairwise correlations within groups (engram, engram-to-be, and other nonengram) computed through the normal model parameters and sleep plasticity OFF model. n = 5, statistical comparisons were made between the normal model and sleep plasticity OFF model using two-way repeated-measures ANOVA, F (1, 4) = 25.57, p = 0.0072 (model data vs sleep plasticity OFF), Sidak’s multiple comparisons test, p = 0.6081 (Engram), p = 0.0001 (engram-to-be), p = 0.9933 (other nonengram). (C) Normalized coincidence ratio during postlearning sleep in the sleep plasticity OFF model. N = 5, statistical comparisons were made between common engram and engram-to-be, specific engram and engram-to-be, common engram and other nonengram, and specific engram and other nonengram, using one-way ANOVA, F (3, 12) = 49.9, p < 0.0001, Dunnett’s multiple comparisons test, p < 0.0378 (common engram and engram-to-be vs specific engram and engram-to-be), p < 0.0001 (common engram and engram-to-be vs common engram and other nonengram), p < 0.0001 (common engram and engram-to-be vs specific engram and other nonengram). (D) Differences in the coincidence ratios between common engram and engram-to-be, and specific engram and engram-to-be, in the normal parameters model and sleep plasticity OFF model. Statistical comparisons were made using a two-tailed t test, t = 5.385, p = 0.007. ^***^p < 0.001, ^****^p < 0.0001, n.s., nonsignificant. Data represent the mean ± s.e.m.

To check if engram-to-be cells show postlearning reactivation, we recorded NREM and REM sleep sessions after session B (Fig. 3F). By implementing population vector distances (PVD), engram-to-be cells showed smaller distances in both NREM and REM sleep compared to other nonengram cells, using session B as a reference. This activity could be representing engram-to-be offline reactivations or replay of session B and behaving similarly as engram cells^8^. To recapitulate, the engram-to-be population behaved in a similar manner during contextual learning for session B but not A’, reactivating at subsequent sleep sessions as a form of consolidation. These implications may designate that this population is the upcoming engram or “engram-to-be” cells for session B.

### Engram and engram-to-be cells show high cooccurrences during postlearning sleep

We next investigated whether there was any interaction between engram cells representing session A and engram-to-be cells representing session B in terms of the coactivity between the two populations within a certain time window. We first tracked the activity of all engram ensembles throughout the sessions. Some ensembles of session A engram cells showed high cosine similarity (>0.6) with session B (common engram ensemble), whereas some ensembles did not (<0.6; (specific engram ensemble, Suppl. Fig. 9A and F). Accordingly, the engram cells were subcategorized according to their activity in session (B) into common engram and specific engram cells (see Methods, Suppl. Fig. 10). To investigate whether there was any functional difference between the two subgroups of the engram population, the neuronal activities were binarized, and all co-occurrences between the different subgroups were counted over a short time window (see Methods; Fig. 4A). The co-occurrences between common engram and engram-to-be cells, and between specific engram and engram-to-be cells, were measured in sessions N1&R1, A, N2&R2, and B (Fig. 4B). Comparisons revealed that the co-occurrences between common engram and engram-to-be cells were significantly higher than the co-occurrences between specific engram and engram-to-be cells only during the postlearning sleep (N2&R2) session, with there being no significant difference detected in the other sessions (Fig. 4C and Suppl. Fig. 10B-D). The differences in the co-occurrences between the subgroups were not due to a difference in the rate of single occurrences of either the common engram or specific cells (Fig. 4D), nor the number of cells (Fig. 4E) in each subgroup. Moreover, there was no significant difference in comparisons of the co-occurrences between common engrams vs. other nonengram cells, and specific engrams vs. other nonengram cells (Fig. 4F). This excludes that the difference in co-occurrences detected during post learning sleep is due to any individual differences of each subgroup in terms of group size or basal activity level. Each of the common and specific engram population represented around 30% of the total engram population (Supp. Fig. 10A). Taken together, these results demonstrate that different subgroups in a single engram population might play different functions in the processing of memory.

### Synaptic plasticity mechanisms during offline periods govern engram-to-be cells emergence

To understand the plasticity mechanism behind the temporal shift of engram assemblies, we created a neural network model to simulate how the engram-to-be cells might emerge. Specifically, we hypothesized that synaptic plasticity during offline periods supports the emergence of engram-to-be cells. In this model, we simulated the responses of CA1 neurons receiving context-dependent inputs from CA3 during four sessions: prelearning sleep, context A, postlearning sleep, and context B (Fig. 5A). During the context exposure, highly activated CA1 neurons become engram cells, and synapses connected to engram cells are potentiated^11,43,44^. With this learning rule, pre-existing assemblies that are active before learning are prone to become engram cells, and after learning, they strongly respond to input patterns of context A (Fig. 5B). During the postlearning sleep, we applied offline synaptic plasticity mechanisms that have been experimentally observed in CA1: synaptic depression in nonengram cells caused by sharp-wave ripples (SWRs)^26^ and homeostatic synaptic scaling^45-47^. After this offline synaptic tuning, nonengram cells in CA1 stopped responding to CA3 patterns of context A, which are frequently replayed during SWRs. Instead, the nonengram cells respond strongly to different CA3 patterns (nonreplay patterns), which results in the emergence of novel assemblies in CA1 (Fig. 5B). These novel assemblies are supposed to be recruited as engram cells if the next experience is different from the previous one, and part of these assemblies are observed as engram-to-be cells. Actually, a previous report showed that such synaptic depotentiation and synaptic scaling during sleep is critical for learning of future memories^26^.

We checked whether the proposed model captures the key features of the experimental data through statistical analyses of simulation results. First, we classified simulated CA1 neurons into engram cells and nonengram cells according to context A, and then nonengram cells were further subdivided into engram-to-be and other nonengram cells based on their activity in context B (see Methods). We subdivided the engram cells of context A into common engram and specific engram types according to their activity in context B (Suppl. Fig.11A; see Methods). The ratio of the cell types in our model matched that in the experimental data (Suppl. Fig. 11B). We analyzed the matching between activity patterns in context A and those in pre and postlearning sleep sessions, and found that the engram cells showed higher matching with context A than the nonengram cells (Fig. 5C), whose matching scores resembled those observed in our in-vivo Ca^2+^ recordings (Fig. 1D). We repeated the same analysis for populations of engram and engram-to-be cells in context B and compared the score against other nonengram cells. The activity patterns of engram and engram-to-be cells showed high matching (compared with other nonengram cells) with context B only during postlearning sleep (Fig. 5D), indicating that the assembly activated in context B was formed after learning of context A. To provide further evidence of the temporal shift in the activity of the engram-to-be cells, we calculated the average pairwise correlation within each cell type (engram, engram-to-be, and other nonengram) in either pre or postlearning sleep sessions. Engram cells showed higher correlations than nonengram cells in both pre and postlearning sleep sessions (Fig. 5E), mimicking the synchronous correlated activity experimentally observed both before and after learning (Fig. 1E). Engram-to-be cells showed high correlated activity only during postlearning sleep, not in the prelearning sleep session, which further validates the temporal shift in the activity of the nonengram cells and the emergence of the engram-to-be cells only after the first learning (Fig. 5E). Next, following subclassification of the engram cells in the experimental data (Fig. 4B), the coincidences of the activities of common engram cells and engram-to-be cells were found to be significantly higher than those of specific engram and engram-to-be cells, and all other combinations, during the postlearning sleep (Fig. 5F), which matches with the experimental result (Fig. 4C). Overall, these results suggest that the hypothesized plasticity mechanism can explain the experimentally observed reformulation of engram assemblies in CA1.

To confirm that offline plasticity is critical for the emergence of engram-to-be cells, we simulated the network model without offline synaptic plasticity in the postlearning sleep period (synaptic depression caused by SWRs and homeostatic synaptic scaling) (Fig. 5B). This manipulation significantly decreased the number of engram-to-be cells and increased the other nonengram cells, while the number of engram cells remained the same (Fig. 6A), indicating that blocking offline sleep plasticity hindered the emergence of the engram-to-be cells. Moreover, the correlation of the activities of engram-to-be cells in postlearning sleep was significantly reduced compared with normal model conditions, while the correlation of activities of engram and other nonengram cells remained the same (Fig. 6B). Comparisons of the correlations of activities between prelearning and postlearning sleep revealed that engram cells increased significantly, engram-to-be cells decreased significantly, and other nonengram cells remained the same. Thus, blocking the offline sleep plasticity affected the engram-to-be cells in the postlearning sleep (Suppl. Fig. 11C). We also measured the difference in the coincidence of activities between common engram and engram-to-be cells, and between specific engram and engram-to-be cells. There was a significant decrease in the model without offline synaptic plasticity (Fig. 6C and D). These results indicate that the emergence of engram-to-be cells depends on offline plasticity during sleep.

## Discussion

Artificially increasing or decreasing the excitability of certain neurons can bias the memory allocation process toward or against them, respectively, thereby generating engram cells^11^. Over the past decade, a consensus has been steadily building regarding the preexisting configuration of the hippocampal network as basis for memory and cognition in rodents^17-19,48-50^ and recently in humans^51^. The intrinsic dynamic activity occurring in the hippocampus is organized into synchronous assemblies, even with minimal external sensory input^49^. This process might act as a physiological equivalent to the artificially activated subsets of neurons, and act as a building block for the upcoming experience^52^. Unique pattern coactivity and hardwiring of the hippocampal preconfigured network were attributed to their birthdate and neurogenesis^53^. A recent study showed that tagging cells while mice are in their home cage immediately before learning, but not 1 day earlier, is sufficient to diminish fear memory if these cells are inhibited during retrieval^54^. However, because of the technical challenges involved, it is difficult to directly relate the activity of engram cells to preconfigured assemblies in freely behaving mice.

Using an imaging system that combines live Ca^2+^ imaging and engram cell labeling allowed us to preemptively monitor the activity of engram cells even before their emergence. Engram ensembles formed during learning showed high correlations during sleep before learning and continued their correlated activities during postlearning sleep. We termed these ensembles ‘preconfigured’ ensembles, which denotes that the brain prepares new ‘slots’ for the next upcoming task. This is in accordance with previous findings showing that engram cells are picked up from a pool of particularly excitable and connected neurons even before learning^54^. Although initial engram cells might have won the competition to encode the first learning experience, it should be possible to recruit a new group of neurons to represent the upcoming experience. Hippocampal neurons are in a continuous dynamic state in which internally generated representations allow new sets of correlated and properly connected cells to fit the requirements of the near future task^36,55^. During postlearning idling periods reactivations take place to consolidate and stabilize previous events^56^. Our results show that, parallel to consolidation a temporal shift in activity occurs in a subpopulation of nonengram to prepare them to the next future event, which we term engram-to-be. Engram-to-be cells developed a temporal shift from a random to a highly correlated pattern of activity by the second learning event during session B, indicating that they were still immature during session A on the first day. The synaptic plasticity mechanisms that occur during idling periods maybe the underlying mechanism for the emergence of these engram-to-be cells. The synchronous activity of engram-to-be cells during new learning signifies the continuous pattern turnover and dynamism in the hippocampal repertoire for future scenarios while the original engram cells preserve the previous memory identity.

Furthermore, we show that engram cells can be further apportioned into two subpopulations: common and specific engrams, each playing a different role. Common engram cells could correspond to general features that can be shared in the future with similar situations, while specific engram cells are tuned to the precise features of the experience that govern its uniqueness. This is in line with previous reports suggesting that within a single engram population there are several subensembles that could represent different features^8^ or have different functions^57,58^.

Sleep and its role in consolidating acquired memories is thoroughly demonstrated^22,24,25^. Both NREM and REM sleep showed comparable results regarding the reappearance of ensembles across sessions. This resemblance in their function could be due to the fact that NREM and REM sleep share several roles in memory processing, such as replay for consolidation^24,25^, or the fact that only a small proportion of the entire sleep was recorded to prevent photobleaching of cells. Majority of engram ensembles were preconfigured aligned, showing high correlated activity from prelearning till postlearning and retrieval periods, whereas stand-by ensembles, were very low (∼5% in REM and 0% in NREM). This indicates that most engram cell ensembles that were active before learning were integrated into the upcoming engram population and maintained their activity after learning for the consolidation process. Sleep is also believed to be of importance for future events^23^. In our network model, we showed that synaptic plasticity in postlearning sleep is crucial for the emergence of engram-to-be that are important for the upcoming event, and also guides the coactivity between engram-to-be and common engram cells. This coupling of the two populations may enhance the basal activity of the engram-to-be cells. This is in agreement with a previous study that showed that optically increasing the excitability of a group of cells unmasks preexisting cell assemblies that were initially non-place cells^19^. Conceptually, this is similar to our engram-to-be cells that were initially masked within the pool of non-engram cells but after the initial learning emerged through synaptic plasticity mechanisms. Collectively, during offline periods and idling moments, two processes occur, consolidating previous memories and preparing for future ones.

Our neural network model not only explains how offline plasticity mechanisms support the emergence of engram-to-be cells, but also suggests an important functional role of the offline state of the brain. Because highly activated cells tend to become engram cells, and engram cells become more excitable^11,43^, the differentiation of engram assemblies for independent memories would require active rebalancing of activity levels. In the offline state, the hippocampal neural network needs to undergo reformulation to prepare novel assemblies for the novel experience while keeping previous engram assemblies stable. Our model provides a specific learning mechanism for such offline tradeoff between assemblies.

We built a minimal model that incorporates key offline plasticity mechanisms to reproduce the experimental data. However, we cannot exclude the possible contribution of other factors. For example, we implemented homeostatic synaptic scaling in our model but tuning of the excitability of each neuron^11,43^ may substitute for this scaling effect. Furthermore, we took into account the plasticity of excitatory synapses from CA3 to CA1 but did not include plasticity of entorhinal cortex-to-CA1 synapses and inhibitory circuits in CA1^59^. Our model explained the emergence of the coactivity between common engram and engram-to-be cells mainly. It is also possible that the common engram cells directly recruit engram-to-be cells through internal connections in CA1. Previous reports suggest that the internal circuit structure in CA1 significantly affects the formation of place coding^60,61^, and that the deactivation of preexisting engram cells results in the rapid recruitment of new engram cells, possibly through internal circuits in CA1^11^. These multiple mechanisms may work independently or cooperatively.

We propose that during offline periods after learning, two simultaneous processes commence: consolidation of past memories through reactivation of engram ensembles, and emergence of an engram-to-be population through synaptic mechanisms to act as a encoders for future memories (Supp. Fig. 12). Together, offline period dynamics fix the past and prepare for the future.

## Author contributions

K.G., N.O., T.F., and K.I. designed the study. N.O. prepared the lentiviruses. K.G. performed the behavioral experiments. K.G. and H.A analyzed the Ca^2+^ imaging data. K.G. and M.R.F performed animal surgery. M.S. established the Thy1-G-CaMP7 transgenic mice. C.C.A.F. performed the mathematical analyses. T.H. constructed the neural model and performed its manipulation. M.N. wrote the code to calculate coactivities and the PCA-ICA analysis. K.G., N.O., T.H., T.F., and K.I. wrote the manuscript. K.I. supervised the entire project.

## Acknowledgments

The authors thank S. Tsujimura and M. Matsuo for genotyping the transgenic mice, Y. Saitoh for helping in the mice surgeries, M. H. Aly for helping in Ca^2+^ imaging data analysis, T. Takekawa for helping with the automatic calcium sorting system (HOTARU), K. Abdou and A. Choucry for their valuable discussion and advice, and all members of the Inokuchi laboratory at the University of Toyama for advice and technical support.

## Data and materials availability

The materials and code that support the findings of this study are available from the corresponding author upon reasonable request. The custom-made program for NMF analysis can be found at https://github.com/fccaa/NMF_custom The custom-made codes for neural network model can be found at https://github.com/oist-ncbc/engramtobe_simulation The custom-made codes for co-activities detection and PCA-ICA analysis can be found at https://github.com/IdlingBrainUT/Ghandour2024_NatureCommunications

## Methods

### Animals

All procedures involving animals complied with the guidelines of the National Institutes of Health, were approved by the Animal Care and Use Committee of the University of Toyama (Approval number: A2022MED-8), and were conducted in accordance with the Institutional Animal Experiment Handling Rules of the University of Toyama. Thy1-G-CaMP7-T2A-DsRed2 mice have been described previously^62^. All surgery was conducted on male c-fos-tTA × Thy1-G-CaMP7 mice with a C57BL/6J background. Mice were maintained on food containing 40 mg/kg doxycycline (Dox) after weaning.

The progeny of the c-fos-tTA × Thy1-G-CaMP7 line were generated by in-vitro fertilization (IVF) of eggs from C57BL/6J mice and embryo transfer techniques to produce a sufficient number of mice for behavioral analysis^3,8^. Genotyping of genomic DNA isolated from the tails of the pups was performed by polymerase chain reaction^63,64^. All mice were maintained on a 12 hour light-dark cycle (lights on at 8:00 am) at 24°C ± 3°C and 55% ± 5% humidity, given food and water ad libitum, and housed with littermates until 5 days before surgery.

### Viral vectors

The pLenti-TRE-hKiKGR plasmid has been described previously^8^. pLenti-TRE-hKiKGR plasmid was prepared using an EndoFree Plasmid Maxi kit (Qiagen). The lentivirus (LV) was prepared as described previously^3^, according to the protocol adopted from that of the laboratory of K. Deisseroth (http://www.stanford.edu/group/dlab/optogenetics/expression_systems.html).

### Stereotactic surgery for Ca^2+^ imaging and EEG/EMG recording

All surgery was conducted on male c-fos-tTA × Thy1-G-CaMP7 mice (aged around 12–20 weeks) with a C57BL/6J background, as described previously ^8^. Briefly, mice were anesthetized by intraperitoneal (i.p.) injection of pentobarbital solution (80 mg/kg of body weight) and placed in a stereotactic apparatus (Narishige, Japan). The craniotomy for injecting TRE-hKikGR lentivirus (LV) was ∼1.0 mm in diameter. TRE-hKiKGR-LV (1 μl/injection site) was injected using an injection cannula connected to a Hamilton microsyringe through a water-filled polyethylene tube. The injection cannula (Unique Medical Co., Ltd., Japan) was targeted to the right CA1 (AP -2.0 mm, ML 1.4 mm, DV -1.5 mm from bregma). LV was injected at 0.1 μl/minute with a microsyringe pump (CMA 400, Harvard Apparatus). After injection, the injection cannula was kept in position for 10 minutes and then slowly withdrawn to prevent any damage to the tissues. The craniotomy was then sealed with dental cement, and the skin was closed.

To ensure complete recovery of the mice, the next surgery to implant a gradient refractive index (GRIN) relay lens over the hippocampus, as reported previously^8,38,65,66^, was performed 4 weeks after the LV injection surgery. Similarly, mice were anesthetized by i.p. injection of pentobarbital solution (80 mg/kg of body weight) and placed in a stereotactic apparatus (Narishige, Japan). A 2.0-mm-diameter craniotomy was made in the skull to set the cannula lens sleeve (1.8 mm outer diameter and 3.6 mm in length; Inscopix, CA). To set the lens over the hippocampus, a cylindrical column of neocortex and corpus callosum above the alveus covering the dorsal hippocampus was carefully and gently aspirated with saline using a 27 gauge blunt drawing-up needle. Phosphate buffered saline (PBS) solution was added frequently to prevent tissue drying up. The cannula lens sleeve was gently placed on the alveus. To fix and straighten the cannula lens sleeve to the edge of the burr hole, bone wax was melted, and the sleeve lens was straightened using low-temperature cautery. The cannula lens sleeve targeted the center of the right hemisphere (AP, 2.0 mm, ML, 1.5 mm). At the frontal part of the skull, two anchor screws linked to a recording connector were fixed to serve as an anchor to the sleeve lens and electrodes for recording the cortical electroencephalogram (EEG). Then, stainless wire electrodes were inserted into the neck muscle to record the electromyogram (EMG). Another set of anchor screws was fixed on the lateral part to act as a body earth for noise reduction. A final set of two screws was fixed on the other side of the skull as an extra anchor for the whole setting. Finally, the visible parts of the skull and electrodes were covered with dental cement to attach the cannula lens sleeve to the skull and anchor screws.

Two weeks after from the sleeve lens and EEG and EMG electrodes setting, mice were anesthetized with pentobarbital solution (80 mg/kg of body weight; intraperitoneal injection), and a GRIN lens (1.0 mm outer diameter and 4.0 mm length; Inscopix, CA) was inserted into the fixed cannula lens sleeve and fixed in place with ultraviolet-curing adhesive (Norland, NOA 81). An integrated miniature microscope (nVista HD, Inscopix, CA)^38^ with a microscope baseplate (Inscopix, CA) was placed above the GRIN lens, allowing observation of G-CaMP7 fluorescence and blood vessels in CA1. The microscope baseplate was fixed to the head of the previously fixed anchor screws using dental cement so that the GRIN lens was covered, and then the integrated microscope was detached from the baseplate. The GRIN lens was covered by attaching the microscope baseplate cover to the baseplate until it was time for Ca^2+^ imaging.

### Microscopy and EEG/EMG recording of freely moving mice

All behavioral analyses and recordings were performed during the light cycle. Ca^2+^ transients were recorded using nVista HD acquisition software (Inscopix, CA) at 20 frames/s frequency, maximum gain, and 60% LED power. To differentiate between different sleep stages, EEG/EMG traces at a sampling rate of 1 kHz with bandpass filtering (low cut-off frequency of 0.5 Hz and high cut-off frequency of 300 Hz for EEG; low cut-off frequency of 5 Hz and high cut-off frequency of 3000 Hz for EMG) were amplified and collected using a Unique Acquisition system (Unique Medical). Using fast Fourier transform analysis, sleep scoring was performed on the EEG and EMG traces for all 16 s consecutive epochs. In this scoring, high EMG and low EEG voltages with high frequency were labeled as wakefulness, low EMG and high EEG voltages with dominant δ (0.5– 4 Hz) frequency components were labeled as non-rapid-eye-movement (NREM) sleep, and low EMG (indicating muscle atonia) and low EEG voltages with dominant θ (6–9 Hz) frequency components were labeled as REM sleep^8^. The contexts used for learning session A and session B were located in the same behavioral room but in different locations. Context A was a cylindrical chamber (300 mm diameter, 450 mm height) with white acrylic walls and a floor covered with brown tissues. Context B was a square chamber with a Plexiglass front, gray sides, back walls (width × depth × height: 175 × 165 × 300 mm), and a floor consisting of 26 stainless steel rods (diameter, 2 mm) placed 5 mm apart. Before starting the experiment, the c-fos-tTA × Thy1-G-CaMP7 mice injected with TRE-KikGR-LV were moved to an isolation box (FRP BIO2000, CLEA Japan) located in a Faraday cage (Narishige, Japan). Mice were first habituated to the weight of the miniature microscope and the EEG/EMG wires for 30 minutes on each of 3 consecutive days. On the next day, DOX pellets were removed from the food and replaced with normal pellets. Two days later, the mice were anesthetized with around 2% isoflurane for about 5 minutes. Then, 15 second pulses of 365 nm light were delivered to the GRIN lens from an LED light source (BL-LED-365, OPTO-LINE, Inc., Japan) through an optic fiber (SH4001 Super Eska, NA: 0.5, Mitsubishi Rayon Co., Ltd., Japan). This procedure was repeated three times with 1-minute intervals between each repetition, to remove any nonspecific KikGR expressed before the learning. Then, a snapshot of the CA1 was captured using the nVista HD acquisition software (Inscopix, CA). Mice with the microscope attached to their head were returned to their home cage in the Faraday cage; the EEG/EMG electrodes were attached, and the mice were allowed to rest. After 1–2 hours, the Ca^2+^ activities of the first stable-appearing NREM sleep were recorded (N1). Around 1 hour later, the first-appearing REM sleep that lasted for at least 1 minute was also recorded (R1). Mice were then introduced to the cylindrical chamber for the first time, and the Ca^2+^ activities were recorded for 6 minutes (session A). After this, the mice were returned to their home cage with the miniature microscope still attached to their head and were allowed to sleep. Around 1 hour later, Ca^2+^ activities of stable NREM (N2) and REM (R2) were recorded, as previously described. The EEG/EMG electrodes and miniature microscope were detached upon waking from the REM sleep. Across the sleep stage recording (N1, R1, N2, and R2), the nVista LED was shut off between NREM and REM for both prelearning and postlearning sleep stages to prevent any photobleaching. Sleep recording duration was recorded for 1 minute for each NREM or REM session and was increased to 3-4 minutes in NREM and 2 minutes REM in a sperate group of mice; both groups showed the same offline reactivation activity (Fig. 1D and Supp. Fig. 4D). The mice were returned to DOX food from the 2^nd^ day. Twenty-four hours after the first context exposure, a snapshot of KikGR expression was captured using the nVista HD acquisition software (Inscopix, CA) under 2% isoflurane anesthesia for 5 minutes, to prevent the capture of any nonspecific spontaneous calcium activity as a KikGR^+^ cell. This was followed by exposure to 15 s pulses of 365 nm light from an LED light source, with each pulse occurring three times with a 1 minute interval between pulses, as mentioned previously. After 1.5–2 hours, Ca^2+^ activities were recorded when the mice were reintroduced into context A for 3 minutes (session A’). Following the re-exposure, the mice with the nVista HD microscope attached were returned immediately to their home cage. Then, after 1.5–2 hours, the mice were introduced into the square context for 3 minutes (session B), and their Ca^2+^ activities were recorded.

In another set of double transgenic mice after injecting the TRE-KikGR LV, setting the EEG/EMG electrodes and fixing the lens and the baseplate for the miniature microscope. Calcium transients were recorded one day earlier (Day 0) before exposure to the learning session during both awake (Aw0) and sleep sessions (S0). Next day (Day 1) both awake (Aw1) and sleep sessions (S1) were captured before session (A) followed by awake (Aw2) and sleep (S2) sessions. Next day (Day 2), KikGR snapshot capturing and photoconversion were performed in the same manner as mentioned earlier, then both awake (Aw3) and sleep sessions (S3) were captured, as well (Supp. Fig. 4C). Later, sleep sessions were recorded (both NREM and REM) after session B (Fig. 3F).

### Ca^2+^ imaging data acquisition, processing, and cell sorting

All Ca^2+^ transients were captured using nVista acquisition software (Inscopix, CA). The captured images were processed in a similar manner to that described in our previous report^8^. Briefly, using Mosaic software (Inscopix, CA), each session movie was spatially binned using a factor of 2. To maintain the same field of view (FOV) and correct for motion artifacts, motion correction was performed (correction type: translation only, spatial mean [r = 20 pixels] subtracted, and spatial mean applied [r = 5 pixels]) using blood vessels as landmarks. The binned movie was then further processed using ImageJ software (NIH). Each session movie was divided (pixel by pixel) by a low-pass filtered (r = 20 pixels) version. Then, the single session movies were concatenated into one total movie containing all the recording sessions to allow tracking of the same neurons across sessions and days. Motion correction was applied to the total movie, using a single frame as a reference frame, to ensure that the XY translation was adjusted across all recording sessions (Supp. Fig. 1A), similar to the method used in a previous report^67^. The motion correction efficiency was assessed by checking the activity of several cells in the total movie across 2 days to ensure that cells maintained both their spatial location and morphology across days (Suppl. Fig. 1B, C, and D). To measure changes in the fluorescence of each frame, the ΔF(t)/F0=(F(t) − F0)/F0 value was calculated for each movie session (where F0 is the mean image obtained by averaging the entire movie for that session). Finally, using Mosaic software, all movie sessions were once again combined into a single movie comprising the entire behavior. Once the Ca^2+^ movie was finalized, a HOTARU fully automated Ca^2+^ sorting algorithm system (high performance optimizer for spike timing and cell location via linear impulse)^40,64^ was used to extract the spatial and temporal dynamics of active cells.

### Identifying engram cells through KikGR expression

Engram cells were identified using the snapshot taken 24 hours after the novel context exposure (session A). The snapshot was first spatially binned by a factor of 2 and then manually realigned to a reference frame to maintain the same FOV, using blood vessels as landmarks. These procedures were performed using Image J software (NIH). Following the procedure used for the Ca^2+^ movies, the snapshot was divided (pixel by pixel) by a low-pass (r = 20 pixels) filtered version. Then, to maintain the top ∼10% signal intensity, a threshold was applied to remove the background. The output image was further processed by binarization (binary options: Iterations, 1; Counts, 2; Close [fill small holes between pixels], and Watershed). Finally, the remaining contours in the resulting image were analyzed automatically using Image J. The successfully detected ROIs represented the locations of KikGR^+^ cells (Suppl. Fig. 2 B-D). Then, the spatial contours generated by the HOTARU system were compared with the spatial locations of the KikGR^+^ cells identified from the Image J ROIs. Some ROIs for gene-expressing cells did not fully overlap with the detected calcium transients and were discarded (Suppl. Fig. 2E). Only ROIs that showed complete overlap with active cells on visual inspection were recognized as engram cells. The engram cells constituted 8% of all automatically detected cells (Suppl. Fig. 2F).

### Selection criteria for engram-to-be cells

Engram-to-be cells were chosen from the nonengram pool of cells by screening all session B ensembles that showed high cosine similarity (>0.6) with any of the postsleep learning (N2, R2) ensembles, while at the same time showing low cosine similarity (<0.6) with all presleep learning (N1, R1). Active neurons participating in these ensembles were further filtered by choosing neurons having activity higher than double the median of the nonzero activity of that ensemble. These selected neurons were later termed engram-to-be cells. Those cells that did not meet these criteria were excluded from the engram-to-be population and added to the other nonengram pool. To identify engram-to-be and other nonengram cells that significantly increased or decreased their activity during session B on day 2, we compared their Ca^2+^ signals for session B with those in session A (Wilcoxon rank-sum test, using a significance threshold of P < 0.01).

### Subcategorization criteria for engram cells

In the screening of the engram ensembles of session A, there were some ensembles that were activated in the prelearning sleep, context A, postlearning sleep, and session B, and other ensembles that were activated in prelearning sleep, context A, and postlearning sleep but not in session B. Engram ensembles that showed activity including session B were termed common engram ensembles, and ensembles that did not show activity in session B were termed specific engram ensembles. The active participating cells (higher than double the median of nonzero activity) of these ensembles were termed common engram and specific engram cells, respectively (Suppl. Fig. 9).

### Ca^2+^ event co-occurrences

The Ca^2+^ transients of each subpopulation of both engram cells (common engram and specific engram) and nonengram cells (engram-to-be and other nonengram) were binarized using a threshold of >3 standard deviations of the ΔF/F signal, and then downsampled from 20 HZ to 4 HZ (250 ms temporal binning) using a custom-made MATLAB code^20^. Consequently, the total co-occurrences between two subpopulations (in a 250 ms time window) were counted for the prelearning sleep (N1&R1), context A, postlearning sleep (N2&R2), and context B sessions (Fig. 4A), and then normalized according to

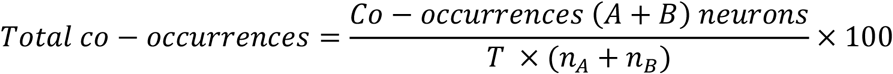

where A is either of the engram subpopulation cells, B is either of the nonengram subpopulation cells, n is the total number of cells active in A or B, and T is the total number of bins. The total co-occurrences were then normalized to the total score of the co-occurrences of specific engrams and engrams-to-be in each calculated session to allow absolute comparisons of co-occurrences across sessions (Fig. 4C). The normalized single occurrences of a certain subpopulation were calculated in a similar manner,

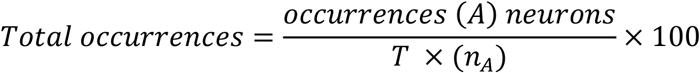

in which A is one of the engram subpopulation cells, n is the total number of cells active in A, and T is the total number of bins. Subjects that did not show at least a single co-occurrence in any or all the sessions were excluded from this analysis (4 of the 9 mice who underwent this analysis were excluded).

### Correlation matrix overlaps

Temporal correlations between pairs of engram cells, engram-to-be cells, other nonengrams, or nonengram cells were calculated using a 1 s sliding time window with time steps of 200 ms. The correlation matrix calculated at time *t* is denoted by 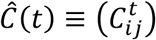, where *i* and *j* are neuron indices, and *t* refers to the midpoint of the time window. Note that if either neuron *i* or *j* was silent, 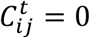. The similarity between the two correlation matrixes at *t* and *t*′ was calculated as 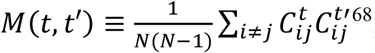, where *N* is the number of cells of interest. To compare engram cells, engram-to-be cells, other nonengram cells, and nonengram cells, the MS with respect to *t*′ was calculated as *M*_total_(*t*) ≡ ∑_*t*′_ *M*(*t, t*′), which measures the degree to which the correlation pattern of the population activity at *t* is repeated at different times within the whole dataset. The larger the value, the more often the correlation pattern looks similar to that obtained at *t*.

Since the numbers of engram cells, engram-to-be cells, other nonengrams, and nonengram cells were different, if the correlation matrixes for different groups of cells were calculated using all cells, the comparison would not be fair. Therefore, we took *N* to be the number of cells in the smallest neuronal group. For those neuronal groups having a size larger than *N*, we took the ensemble average of *M*_total_(*t*) to be the representative sum of the correlation matrixes. Each ensemble consisted of randomly sampled *N* cells from the group.

### Cell ensemble analysis

#### Non-negative matrix factorization (NMF)

NMF was used to extract population activity patterns from the calcium signal dataset^8^. More specifically, NMF finds an optimal factorization of the data matrix 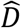, with a pattern matrix 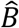 and the corresponding intensity matrix 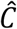, i.e., 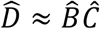. Here, the rows of 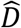 represent the time series of the signals from individual neurons, each column vector of 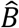 represents a synchronously activated neuron ensemble (population activity pattern), and each row of 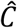 represents the time series of the activation intensity of the corresponding pattern. To search for such a factorization, the cost function defined by *E* ≡ ∑_*ij*_(*D*_*ij*_ − ∑_*k*_ *B*_*ik*_*C*_*kj*_)^2^ was minimized using both multiplicative and additive algorithms^42^. Random initial entries from matrixes 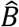 and 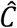 were used for 1000 minimization attempts, and the pair of pattern and intensity matrixes that minimized the cost function were chosen to be the best factorization. The cost function becomes smaller if more patterns are introduced, but the model (i.e., the right-hand side of the cost function) becomes more complex. This trade-off between cost minimization and model complexity was optimized using the Akaike information criterion with second-order correction (AICc)^41^ to determine the optimal number of patterns (i.e., the number of columns in the pattern matrix). Both the neuron indices and the timestamps of the original data were shuffled independently to obtain a shuffled dataset. Ten shuffled datasets were constructed for each sample in each session. Because the ratio of the standard deviation to the mean was small (∼0.03), this number of samples should be sufficient.

#### Matching score (MS)

The overall similarity between pattern vectors in sessions X and Y was measured according to the normalized dot product 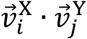 for all possible pattern pairs across the two sessions, noting that the dot product is equivalent to the cosine of the angle between the pattern vectors^69,70^. To this end, the MS between sessions X and Y was defined as 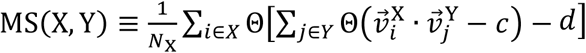, where 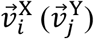 is the *i*-th (*j*-th) pattern vector in session X (Y), *N*_X_ is the number of patterns in session X, and Θ(⋅) is a step function. In supplementary figure 4A, to make the comparison between engram and nonengram cells fair, the number of those major principal components is the same as the number of engram cells in the concerned sample. the MS between sessions X and Y was defined as 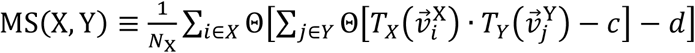, where 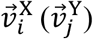 is the *i*-th (*j*-th) pattern vector in session X (Y), *N*_X_ is the number of patterns in session X and Θ(⋅) is a step function. *T*_*X*_ (*T*_*Y*_) is a vector-valued mapping projecting vector 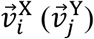 onto major eigen vectors of session X (Y). The constant *d* is an arbitrary positive number smaller than unity. This scoring function yields the portion of patterns in session X that have a normalized dot product larger than *c* with any of the patterns in session Y. A threshold of *c* = 0.6 was used throughout the study. Monte Carlo resampling was conducted for the MSs of engram cells. In the resampling, the MSs of datasets with shuffled timestamps and neuronal indices were calculated to assess significance. For each session-pair comparison, the mean of the MS for the shuffled data was compared with that calculated from the original data. To calculate the Mean of the MS for the shuffled data, 40 shuffled samples were used. Since the resampling was not for the purpose of constructing a random-number distribution, the calculation of the mean value converged, and the number of samples was sufficient. Normalized data were calculated by subtracting shuffled data of engram and nonengram cells from their corresponding MS data.

#### Principal component analysis

To detect coactivity patterns among calcium transients of the total nonengram population (engram-to-be and other nonengram), an unsupervised analysis combining Principal Component Analysis (PCA) and Independent Component Analysis (ICA) was employed, as described previously^71,72^. Briefly, neuronal activities were binned in 20 ms time bins and normalized (z scored, in which the activity of every neuron was adjusted to have an average value of zero and a standard deviation of one) to prevent any bias by neurons having a higher firing rate. We derived ensembles of coactive neurons through a two-step process. Initially, we estimated the quantity of meaningful coactivation ensembles present within a certain population of neurons by identifying the number of principal components from the activity matrix. Subsequently, we employed an independent component analysis to extract the coactivity patterns defined by these principal components (Suppl. Fig. 8), as described in previous reports^71,72^.

### Population Vector Distance (PVD)

PVD quantifies the difference in calcium signals obtained from a group of neurons during two distinct sessions. Calcium signals from two distinct sessions, X and Y, of the same neuron group were compared using the restricted Mahalanobis distance, which is defined on the basis of the Mahalanobis distance^73^. The restricted Mahalanobis distance is defined as 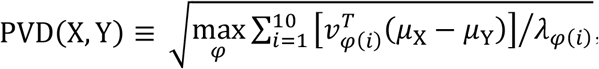, where *μ*_X_ and *μ*_Y_ are the means of X and Y, respectively, *λ*_*φ*(*i*)_s are eigenvalues of the covariance matrix of the union of X and Y, *v*_*φ*(*i*)_ is the corresponding eigenvector, and *φ* is a bijective mapping reordering indices of eigenvalues and eigenvectors. Intuitively, this equation calculates the maximum distance within a restricted number of dimensions. This definition enables us to compare the PVD across different groups of neurons, i.e., engram-to-be cells and other nonengram cells, that contain different numbers of neurons.

### Simulation model and procedure

In our network model, we set the activities of CA3 neurons *r*_*i*_ (*i* = 1,2, …, *N*_CA3_) as inputs, and simulated the responses of excitatory neurons *x*_*i*_ (*i* = 1,2, …, *N*_CA1_) and inhibitory neurons *y*_*i*_ (*i* = 1,2, …, *N*_inh_) in CA1. The simulations were performed by solving dynamical equations:

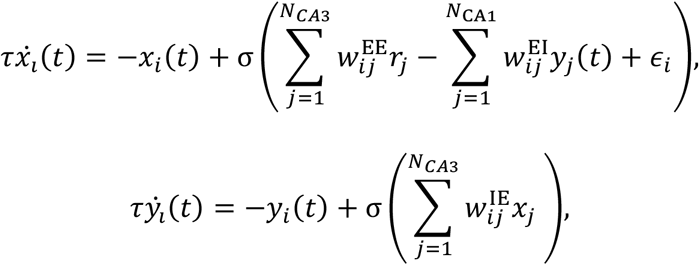

where σ(*x*) is a logistic sigmoid function:

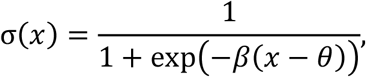

and *ϵ*_*i*_ is a random external input sampled from a Gaussian distribution with a standard deviation of 0.5. The number of neurons was *N*_CA3_ = *N*_CA1_ = 400 and *N*_inh_ = 100. The time constant was *τ* = 2 ms. The parameters of the sigmoid function were *β* = 5 and *θ* = 1. For each CA3 activity pattern, we initialized *x*_*i*_(*t*) = 0 and *y*_*i*_(*t*) = 0 and sampled *ϵ*_*i*_, the simulated time evolution of the dynamics for 20 ms, and recorded the activity patterns of CA1 excitatory neurons at the final state as a response. Below, we specify a CA3 activity pattern as a *N*_CA3_-dimensional vector **r** and a CA1 response pattern as a *N*_CA1_-dimensional vector **x**.

In each CA3 pattern **r**, each neuron took a value of 1 (active) with a 10% probability and was otherwise zero (inactive). In simulations of sleep sessions, the values of active neurons were randomly fluctuated within the range of 0.5–1.5.

Before each simulation, we initially sampled the excitatory synaptic weights 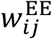 from a uniform distribution [0, 0.125]. We set the I-to-E weight 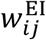 to 1.6 with a 5% probability, with it being zero otherwise, and the E-to-I weight 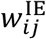 to 0.4 with a 10% probability and zero otherwise.

The simulation consisted of four sessions: presleep, context A, postsleep, and context B. First, we performed a prelearning sleep session in which we recorded CA1 responses 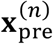 to 1000 random CA3 patterns 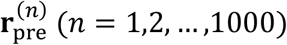. Next, we simulated a CA1 response **x**_A_ in the context A by setting a CA3 pattern **r**_A_ (**r**_A_ was randomly sampled and fixed throughout the simulation). After the first simulation, active CA1 neurons (*x*_*i*_ > 0.5 in **x**_A_) were labeled as engram cells. We defined an engram label 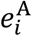 that takes 1 for engram cells and 0 for nonengram cells. Using these labels, we potentiated 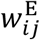 as

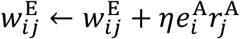

Here, 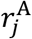 represents CA3 activities in the context A (**r**_A_) and *η* = 0.05. We performed a second simulation using updated weights and recorded a CA1 response **x**_A_ in the second simulation for data analyses (**x**_A_ in the first simulation was discarded).

In the postlearning sleep session, we first applied synaptic depression caused by SWR and synaptic scaling. The synaptic depression caused by SWR is defined by

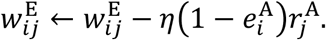

The weight updates resulting from the synaptic scaling were

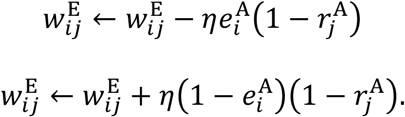

After updating the weights, we obtained CA1 responses 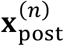 to 1000 CA3 activity patterns 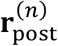.

Eighty percent of 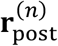 was the same as **r**_A_ (replay), and the remaining 20% were independent random patterns.

Finally, we simulated a CA1 response **x**_B_ in the context B by setting a CA3 pattern **r**_B_. We used the same procedure as used for context A. We determined engram cells for the context B first, potentiated synapses, and recorded a CA1 response **x**_B_ with updated weights.

We repeated this simulation procedure five times with different random seeds and analyzed data from each trial independently. In the control simulations presented in Fig. 6, we did not apply synaptic changes in postsleep sessions (synaptic depression by SWR and synaptic scaling) while keeping synaptic potentiation in context A and B.

### Analysis of simulated data

We separated engram cells and nonengram cells by engram labels determined from the first simulation in context A. Furthermore, we separated active and inactive cells in context B by thresholding the activity (a neuron was active if *x*_*i*_ > 0.5). Accordingly, we defined four subtypes of CA1 neurons: common engram cells (engram in context A and active in context B), specific engram cells (engram in context A and inactive in B), engram-to-be cells (nonengram in context A and active in B), and other nonengram cells (nonengram in context A and inactive in B). The activity patterns below (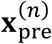, **x**_A_, 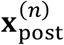, and **x**_B_) indicate divided parts of patterns based on cell types.

The matching ratio between presleep and context A was calculated as

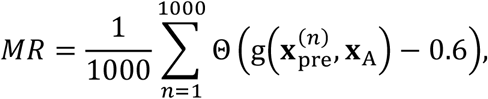

where Θ(*x*) is a Heaviside step function, and g(**x, y**) is the cosine similarity between vectors **x** and **y**. Matching ratios for other combinations (such as postlearning sleep and context B) were also calculated using the same procedure.

To calculate pairwise correlations and coincidence ratios, we artificially generated the augmented CA1 activity patterns 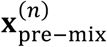 and 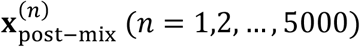 by mixing 1000 simulated activity patterns and 4000 “silent” activity patterns in which the activity of each CA1 neuron was sampled from the range [0, 0.01]. These silent activity patterns corresponded to activity-silent periods between burst activities in CA1. The correlation within each cell type was calculated as follows. The activities of each CA1 neuron in a sleep session were expressed as 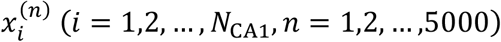. The correlation coefficient between neuron *i* and *j* was calculated using Pearson correlation between two sets of observations 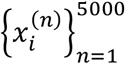 and 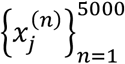.

We averaged the correlation over all cell pairs within each cell type (engram, nonengram, or one of the four subtypes). The coincidence ratio was calculated using the mean population activity in a cell type 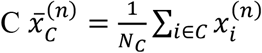 (C is common engram, specific engram, engram-to-be, or other nonengram), where *N*_*C*_ is the number of neurons in cell-type C. Furthermore, to calculate the baselines of the coincidence, we generated shuffled activities 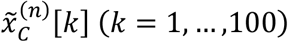 by shuffling the index (n) of 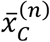 100 times. The coincidence ratio between two cell types C1 and C2 was calculated as

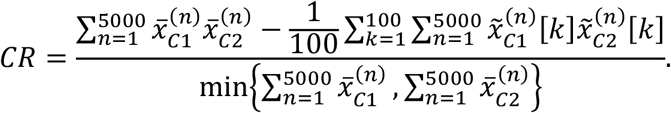

### Statistical analysis

Statistical analyses were performed in GraphPad Prism 6 (GraphPad Software). Comparisons of data between three groups were performed using one-way ANOVA. Comparisons of data between two groups at multiple time points were performed using two-way repeated-measures ANOVA followed by Bonferroni’s multiple comparisons test, unless otherwise specified. Comparisons of data between two paired groups were performed using paired *t*-tests (no assumption was made that data were normally distributed). One-tailed comparisons were used whenever the difference between the two groups was expected to be in a single direction. Quantitative data are expressed as the mean ± standard error of the mean (s.e.m.).

## Supplementary Information

**Suppl. Fig.1.**
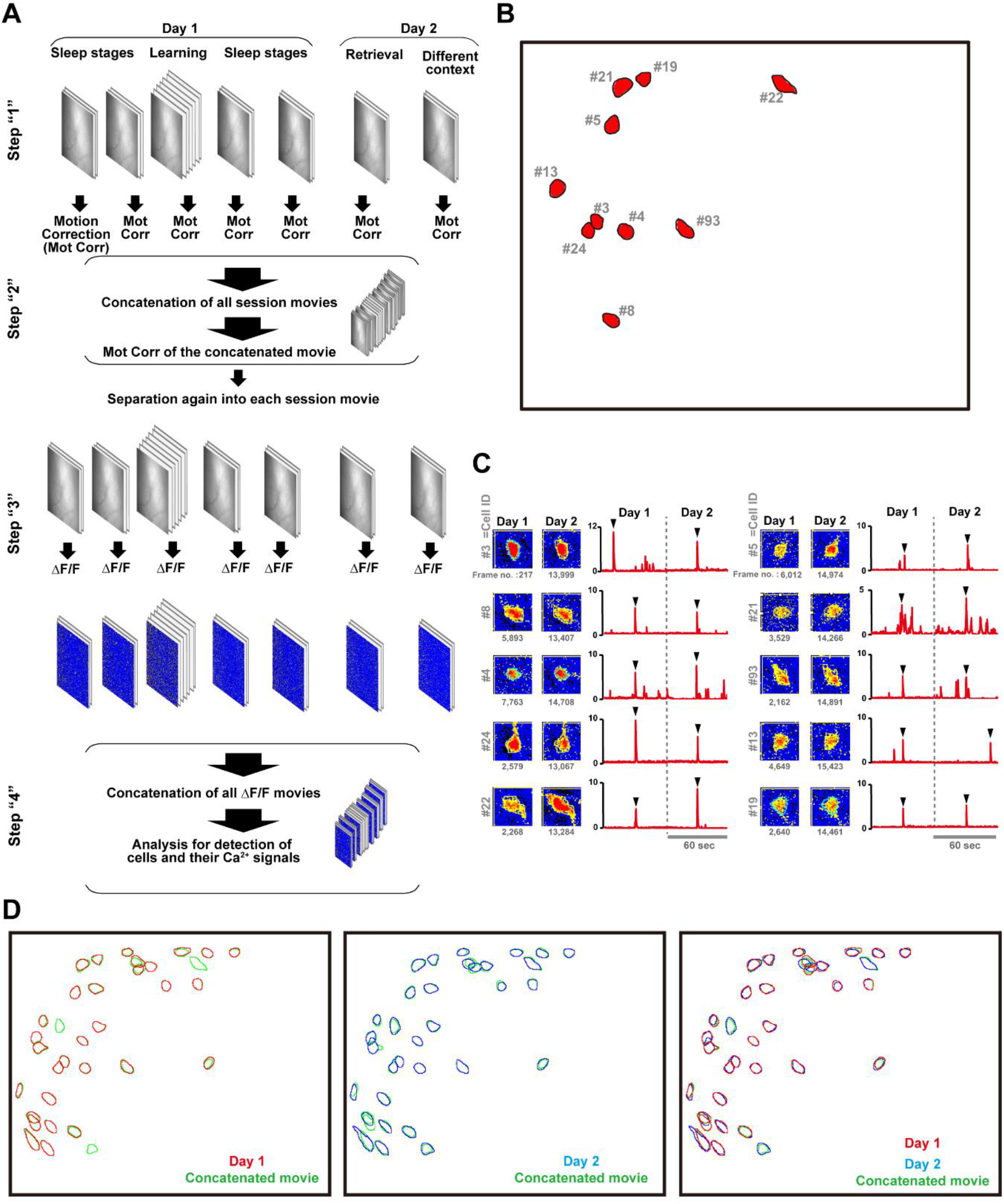
Extracting calcium transients from engram cells and nonengram cells across sessions maintaining the same field of view (FOV). (A) Schematic diagram showing the preprocessing of a Ca2+ imaging video of the whole recorded behavioral paradigm. (B) Spatial positions of representative cells in the FOV that were active on both days of recording. (C) ROI of the selected cells in panel (B), and their representative Ca2+ traces on day 1 and day 2. Black arrowheads point to the time selected for evaluating cell morphology. The number of frames is indicated under each ROI. (D) Footprints of matched cells detected by the HOTARU system from the concatenated movie and videos of single days (Days 1 & 2). Cells detected from the concatenated video are shown in green (left, center, right), cells detected from the day 1 video are shown in red (left, right), and cells detected from the day 2 video are shown in blue (center, right).

**Fig.2.**
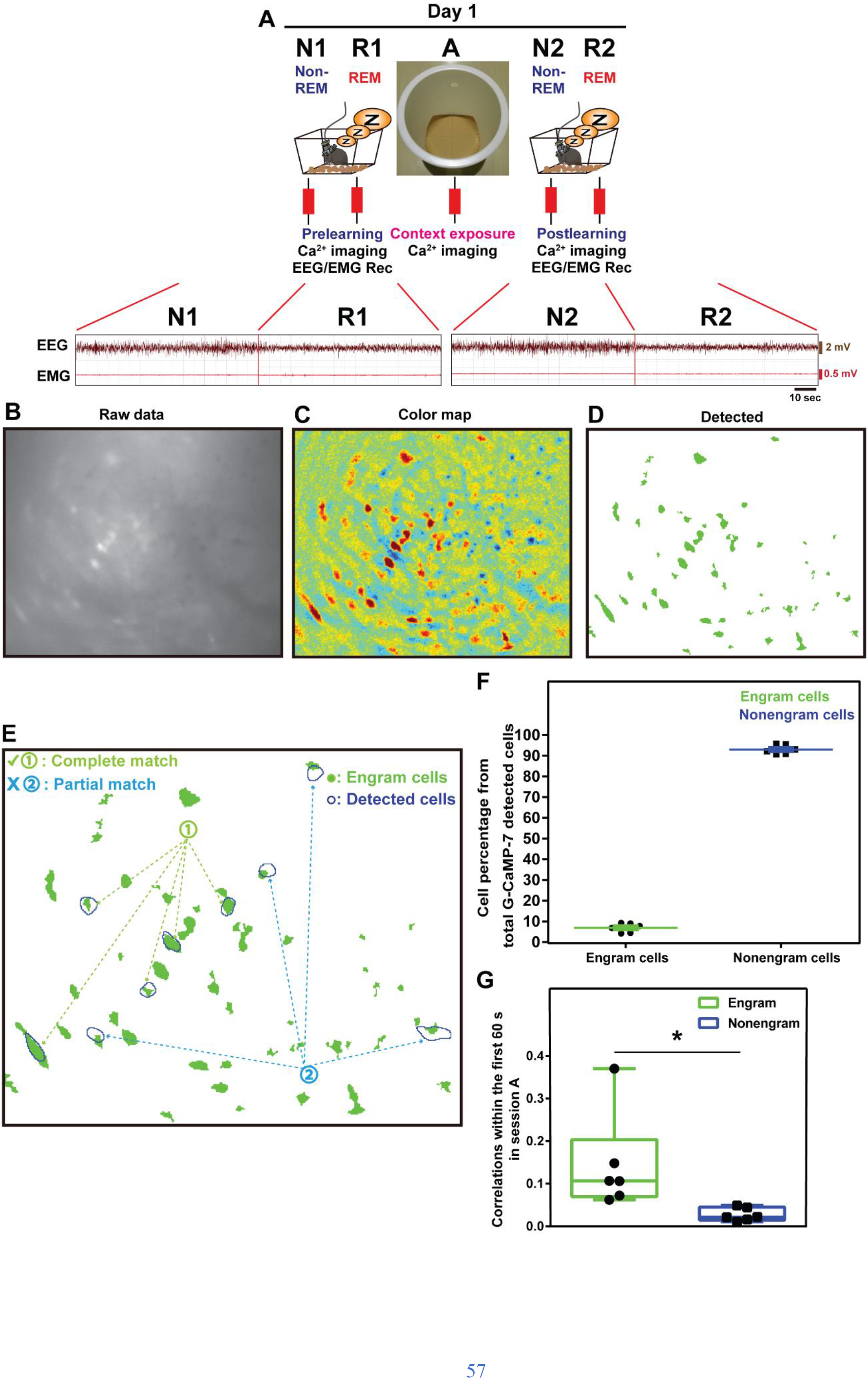
Electroencephalogram (EEG) and electromyogram (EMG) traces and identification of double-positive KikGR and G-CaMP-7 cells. (A) Representative EEG and EMG traces during calcium imaging sleep recording to differentiate between nonrapid eye movement (NREM) and rapid eye movement (REM) sleep stages, in prelearning (N1 and R1) and postlearning (N2 and R2) sleep. (B-D) Representative snapshot image of KikGR+ expression raw data (A), colored map (B), and automatic detection of KikGR+ (C) (see Methods). (E) Criteria for selecting KikGR+ cells. Cells showing a complete match between the KikGR-expressing cell from the snapshot (green filled contours) and the HOTARU system (blue contours; category 1; i.e., cells double-positive for c-fos tet tagging and G-CaMP-7 fluorescence) were considered engram cells. Cells that showed only a partial match between the blue and green contours (category 2) were excluded from further analysis and considered as nonengram cells. (F) Percentage of engram cells (KikGR+ & G-CaMP7+) to nonengram cells (total detected GCaMP7+). (G) Average overlaps in engram cells and non-engram cells in session A, the summation of Mtot(t) over time during the first 60 s of Ca2+ imaging relative to the sum from nonengram cells during novel context exposure (session A). Statistical comparisons were conducted using a two-tailed t test. n = 6; (engram vs nonengram) t (5) = 2.738, p = 0.0409; ^*^p < 0.05. Data represent the median ± s.e.m.

**Suppl. Fig. 3.**
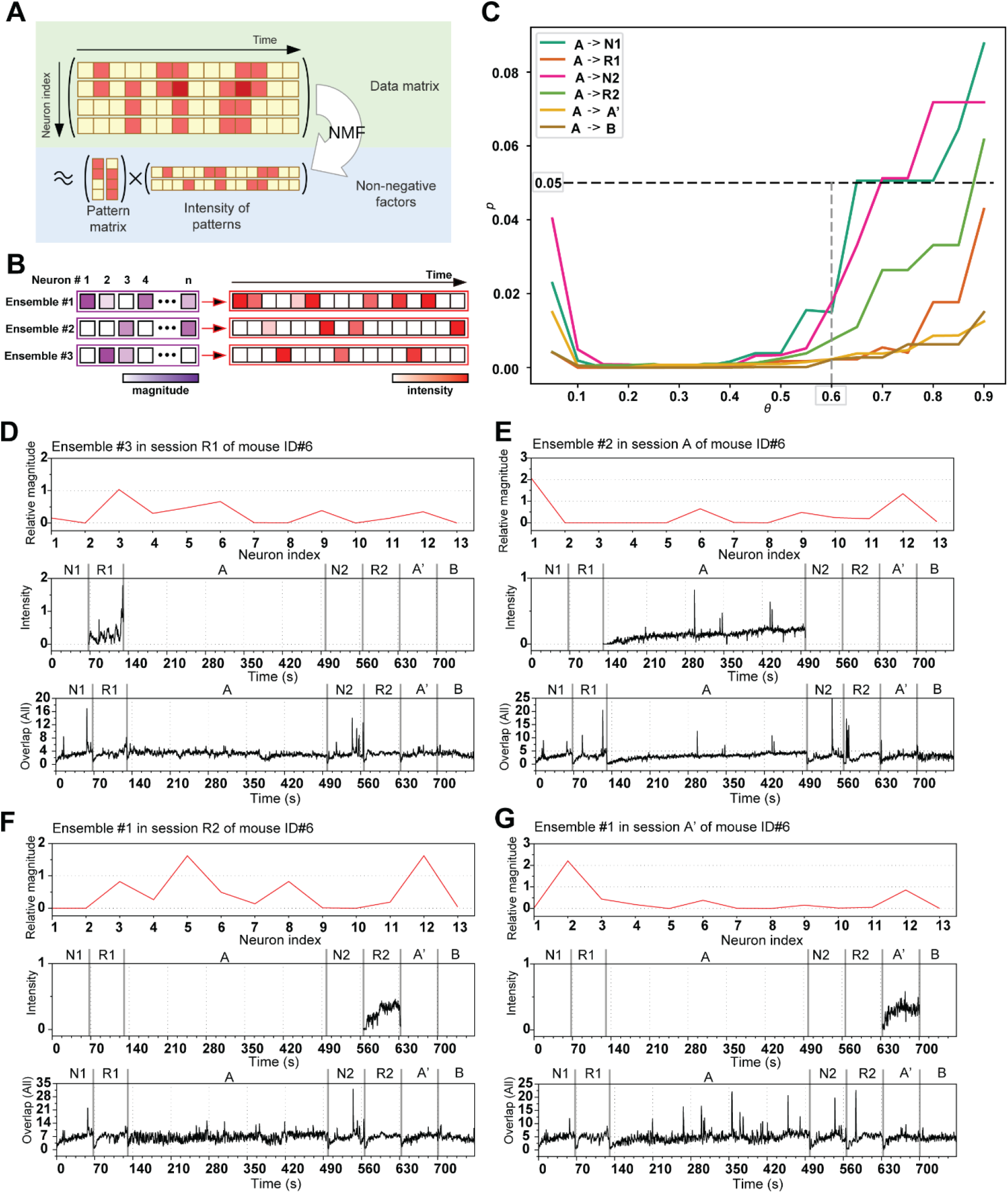
Ensembles detected by NMF analysis across sessions. (A-B) Illustrative diagram of the non-negative matrix factorization (NMF) analysis (A), and the relative magnitude of each neuron participating in that ensemble and the activity of that ensemble across time (B). (C) Summary of *p*-values for the difference between the MSs for engram and non-engram cells in each session (at several dot product thresholds). Using paired t-test between engram and non-engram cells. (D-G) Representative ensembles detected by the NMF analysis showing the neurons participating in that ensemble (top) and their relative intensity in different sessions; R1 (D), A (E), R2 (F), and A’ (G) (middle), and their relative intensities across all the sessions (bottom).

**Supp. Fig. 4.**
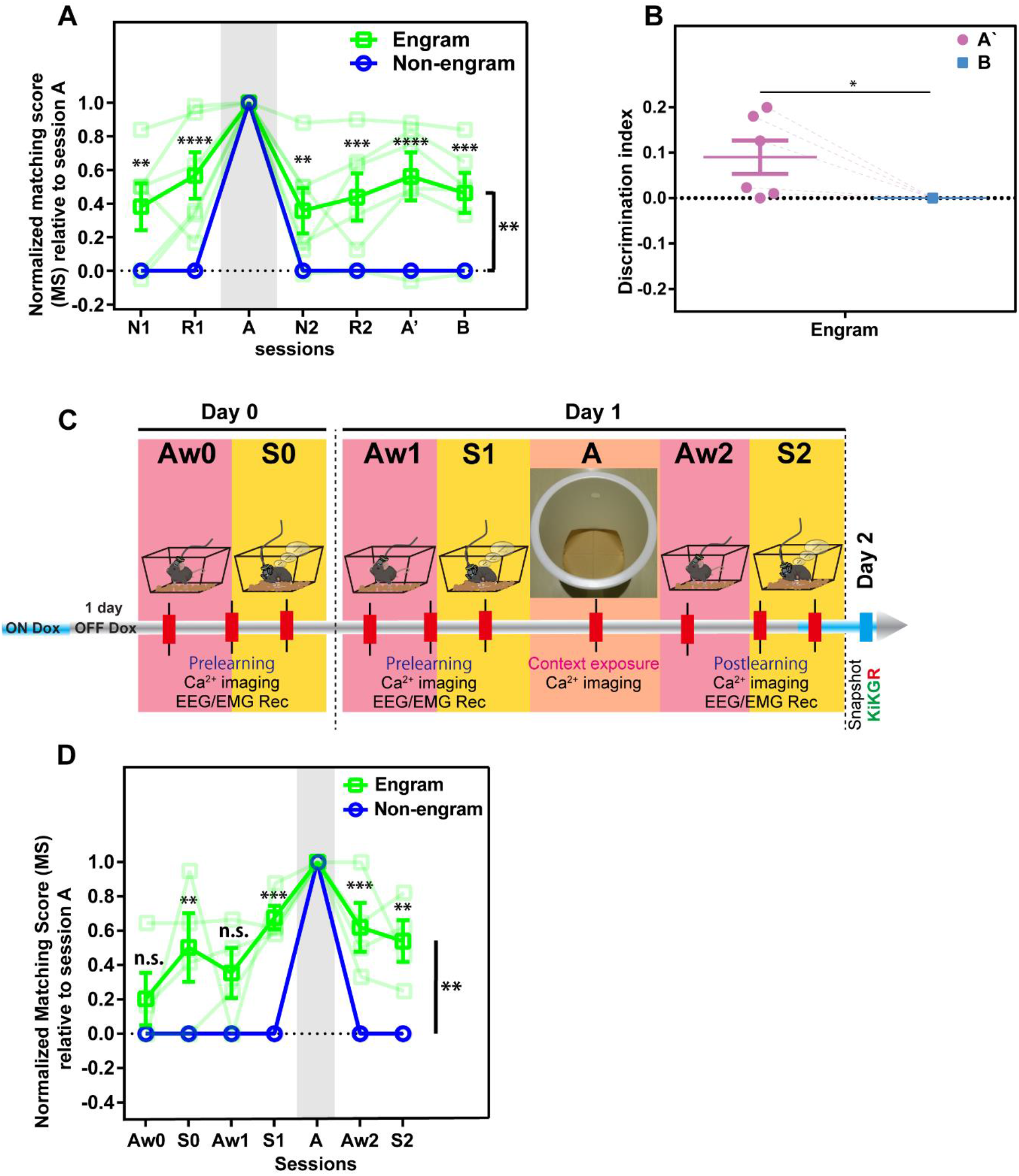
Sleep but not awake activity is more correlated to next learning event. (A) MS analysis in reference to session A (same as Fig.1D but with using same number of neurons to run the analysis). n = 6, statistical comparisons were made between engram and nonengram using two-way repeated-measures ANOVA, F (1, 5) = 21.36, p = 0.0057 (engram vs nonengram), Bonferroni’s multiple comparisons test, p = 0.0043 (N1), p = <0.0001 (R1), p = 0.008 (N2), p = 0.0009 (R2), p = <0.0001 (A’), p = 0.0004 (B). (B) Discrimination index between sessions A’ and B for engram cells, n = 6, statistical comparisons were made between A’ and B using one-way paired t-test, t (5) = 2.405, p = 0.03. (C) Experimental design for calcium imaging and awake, sleep behavior paradigm. Awake (Aw0) and sleep (S0) recorded session at 1 day before learning (Day 0), and then awake (Aw1) and sleep (S1) in prelearning sleep sessions, then learning (A), and followed by postlearning awake (Aw2) and sleep sessions (S2) on day 1. On day 2, a snapshot for KikGR is taken. (D) Normalized MS analysis in reference to session A for both engram (green) and nonengram (blue) cells across sessions. n = 4, statistical comparisons were made between engram and nonengram using two-way repeated-measures ANOVA, F (1, 3) = 37.30, p = 0.0088 (engram vs nonengram), Bonferroni’s multiple comparisons test, p = 0.8289 (Aw0), p = 0.0052 (S0), p = 0.0734 (Aw1), p = 0.0002 (S1), p = 0.0006 (Aw2), p = 0.0027 (S2).

**Suppl. Fig. 5.**
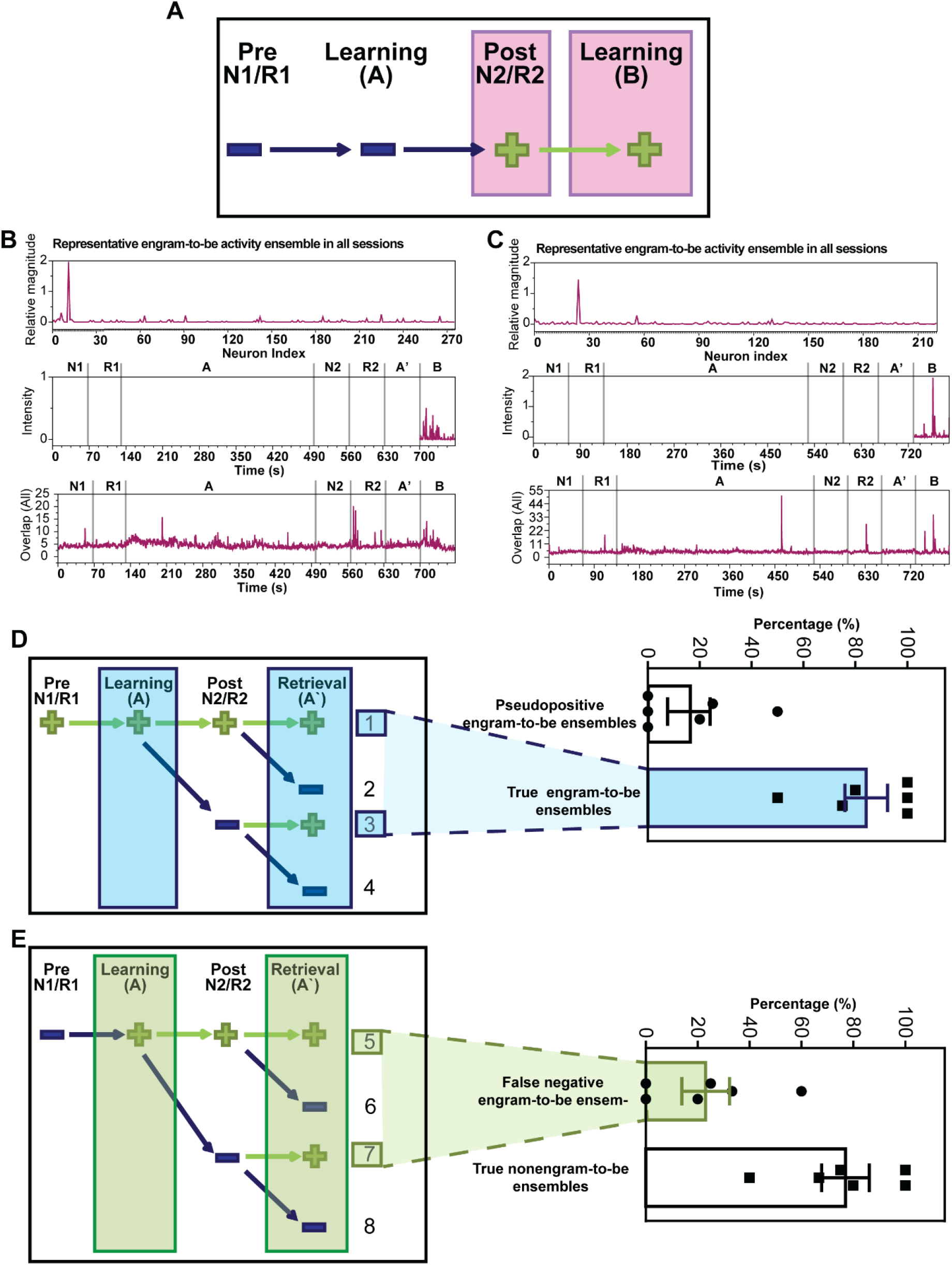
Engram ensemble activity across sessions. (A) Schematic diagram showing engram-to-be ensemble activities across sessions. (B and C) Representative engram-to-be ensembles detected by NMF analysis showing the neurons participating in that ensemble (top), their relative intensity in session B (middle), and their relative intensities across all sessions (bottom). (D) Left, schematic diagram showing all possible engram-to-be ensembles according to the criteria, in a similar manner to Figure 2. Right, percentages of true engram-to-be ensembles that were preactivated and retrieved (patterns 1 and 3), and pseudopositive engram-to-be ensembles that were preactivated and not retrieved (patterns 2 and 4). Ensemble percentages for each category were calculated as follows: (True= (1+3)/ (1+2+3+4)), (Pseudo-positive= (2+4)/ (1+2+3+4)). (E) Left, schematic diagram showing all possible ensembles assigned as non-engram-to-be ensembles. Right, percentage of false-negative engram-to-be ensembles (true engram-to-be ensembles) that were not preactivated but activated during learning and retrieval (patterns 5 and 7), and true nonengram-to-be ensembles not preactivated and not retrieved (patterns 6 and 8). False negative= (5+7)/ (1+3+5+7), True negative= (6+8)/ (1+3+5+7).

**Suppl. Fig. 6.**
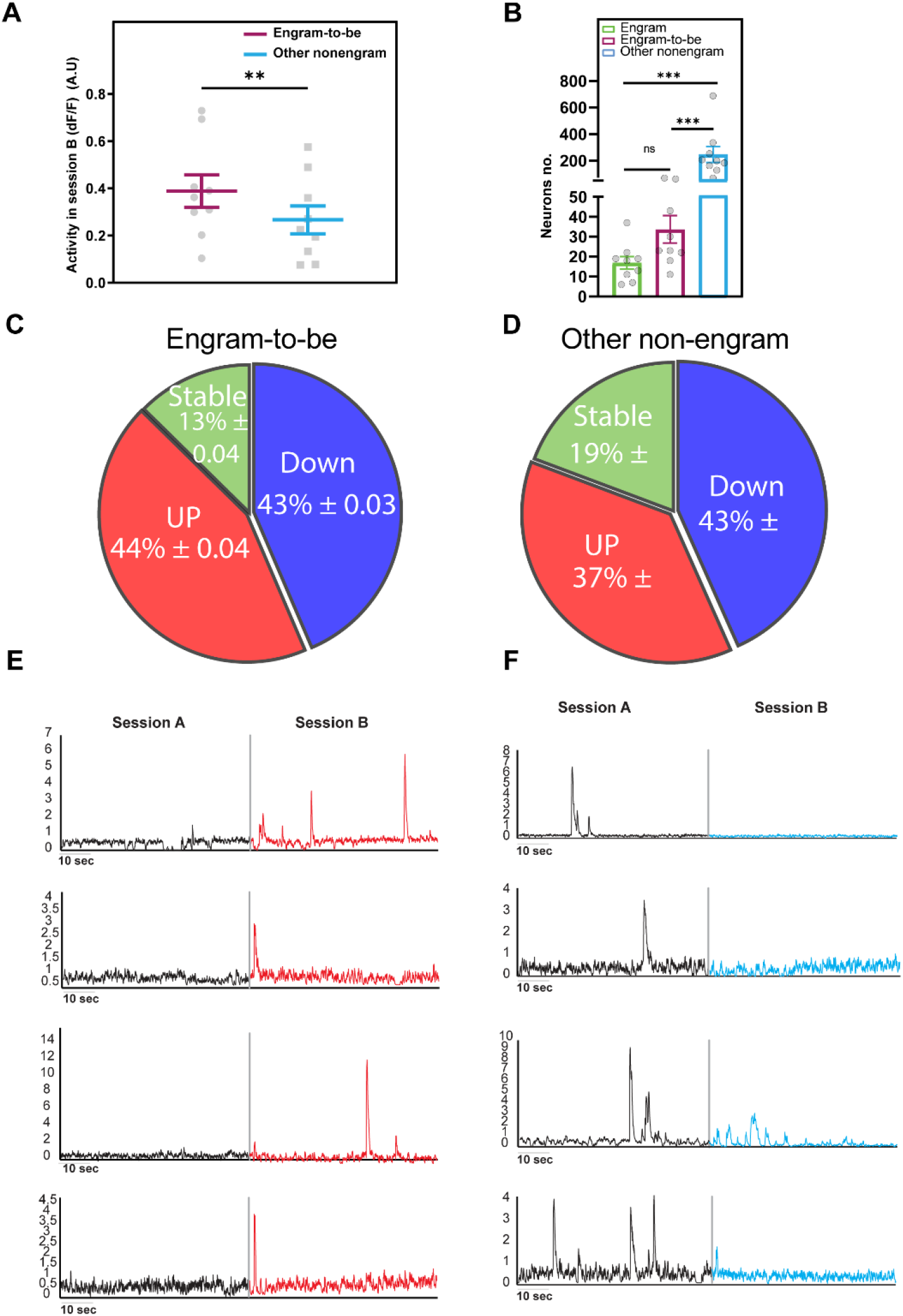
The engram-to-be population showing higher activity in session B than in session A. (A) Basal activity of engram-to-be and other nonengram in session B. n = 9, two-tailed t test between engram-to-be and other nonengram, t (8) =4.843, p = 0.0013. (B) The number of neurons in engram (green), engram-to-be (magenta), and other nonengram cells (cyan). Statistical comparisons were made using one-way ANOVA, n = 9, F = 13.39, p = 0.0001 (between engram, engram-to-be, and other nonengram). Tukey’s multiple comparisons test, p = 0.9393 (engram vs engram-to-be), p = 0.0003 (other nonengram vs engram), p = 0.0007 (other nonengram vs engram-to-be). (C) Percentage of engram-to-be cells showing a significant increase in their activity in session B compared with session A as the up population, cells showing a significant decrease as the down population, and cells that did not show a significant change as the stable population, shown in red, blue, and grey, respectively. (D) Representative traces of engram-to-be cells from the up population showing a significant increase in their activity in session B compared with session A. (E) Percentage of other nonengram cells showing a significant increase in their activity in session B compared with session A as the up population, cells showing a significant decrease as the down population, and cells that did not show a significant change as the stable population, shown in red, blue, and grey, respectively. (F) Representative traces of other nonengram cells from the down population showing a significant decrease in their activity in session B compared with session A.

**Suppl. Fig. 7.**
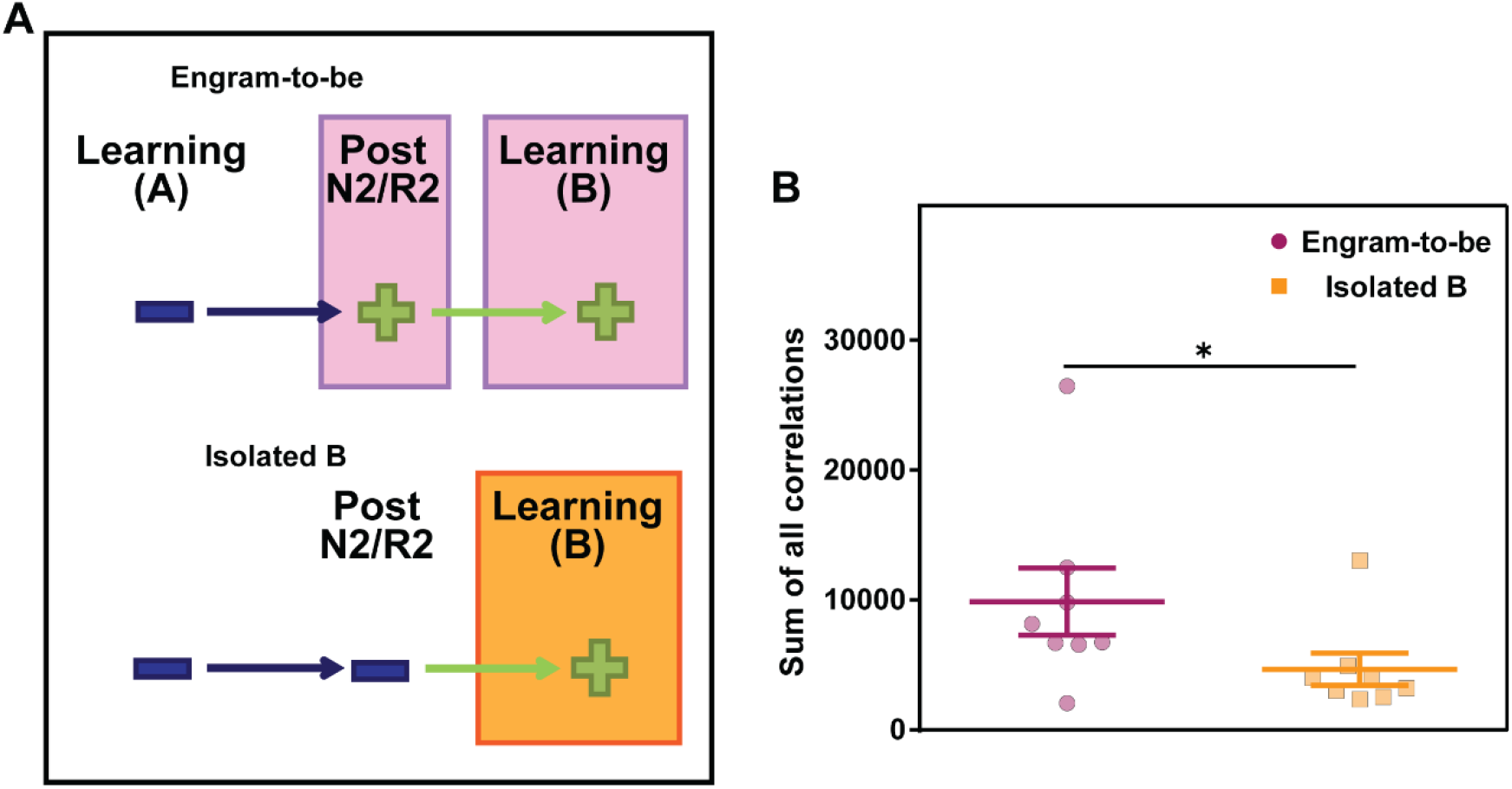
Engram-to-be cells show more correlated activity than isolated B ensembles in session B. (A) Schematic diagram showing engram-to-be (top) and isolated B (bottom) ensemble activities across sessions. (B) Sum of all correlations (Mtot(t)) in the first 60 seconds for engram-to-be and isolated B in session B. Statistical comparisons were made using two-way paired t test, n = 8, t = 3.454, df= 7, p = 0.0106 (engram-to-be vs isolated B), ^*^p < 0.05. Data represents the mean ± s.e.m.

**Suppl. Fig. 8.**
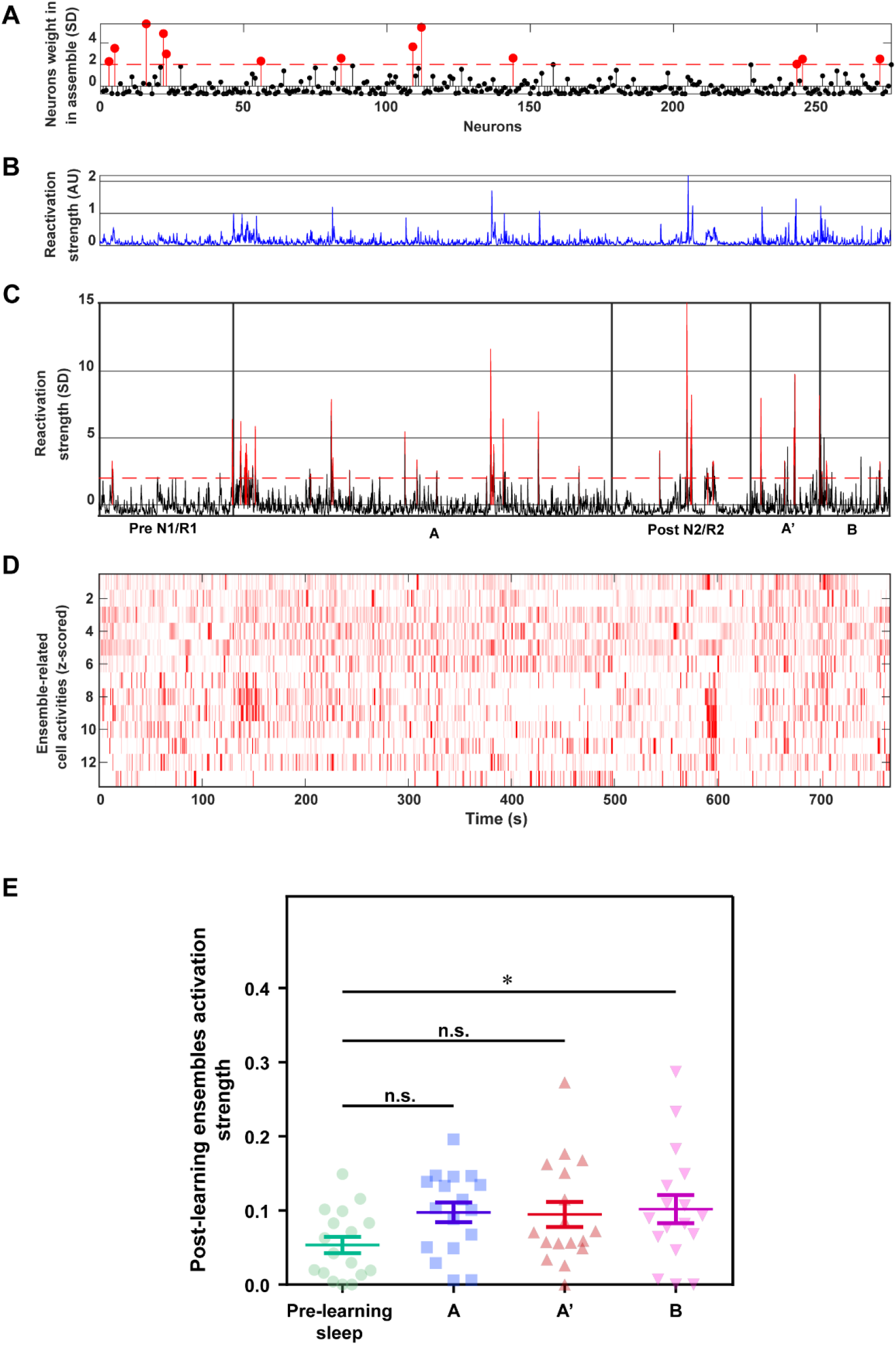
PCA-ICA analysis shows higher activation strength of postlearning sleep ensembles during session B. (A) Weight of neurons participating in a representative ensemble, red dots are significant participating neurons (>2SD), red dashed linerepresents threshold (2SD). (B) Reactivation strength (AU) of a given representative ensemble in sessions prelearning sleep, A, A’, postlearning sleep and B. (C) Reactivation strength (SD) across the whole sessions, where red dashed line indicates threshold for activity (2SD) in sessions prelearning sleep, A, A’, postlearning sleep and B. (D) Activities of the 13 neurons participating in the ensemble (z-score) across time. (E) Activation strength of all detected in ensembles in postlearning sleep in other sessions. Statistical comparisons were made using one-way RM ANOVA, N = 6, n = 17, F = 2.56, p = 0.0662. Dunnett’s multiple comparisons test, p = 0.0495 (prelearning sleep vs B), p = 0.9932 (A vs B), p = 0.9716 (A’ vs B).

**Suppl. Fig. 9.**
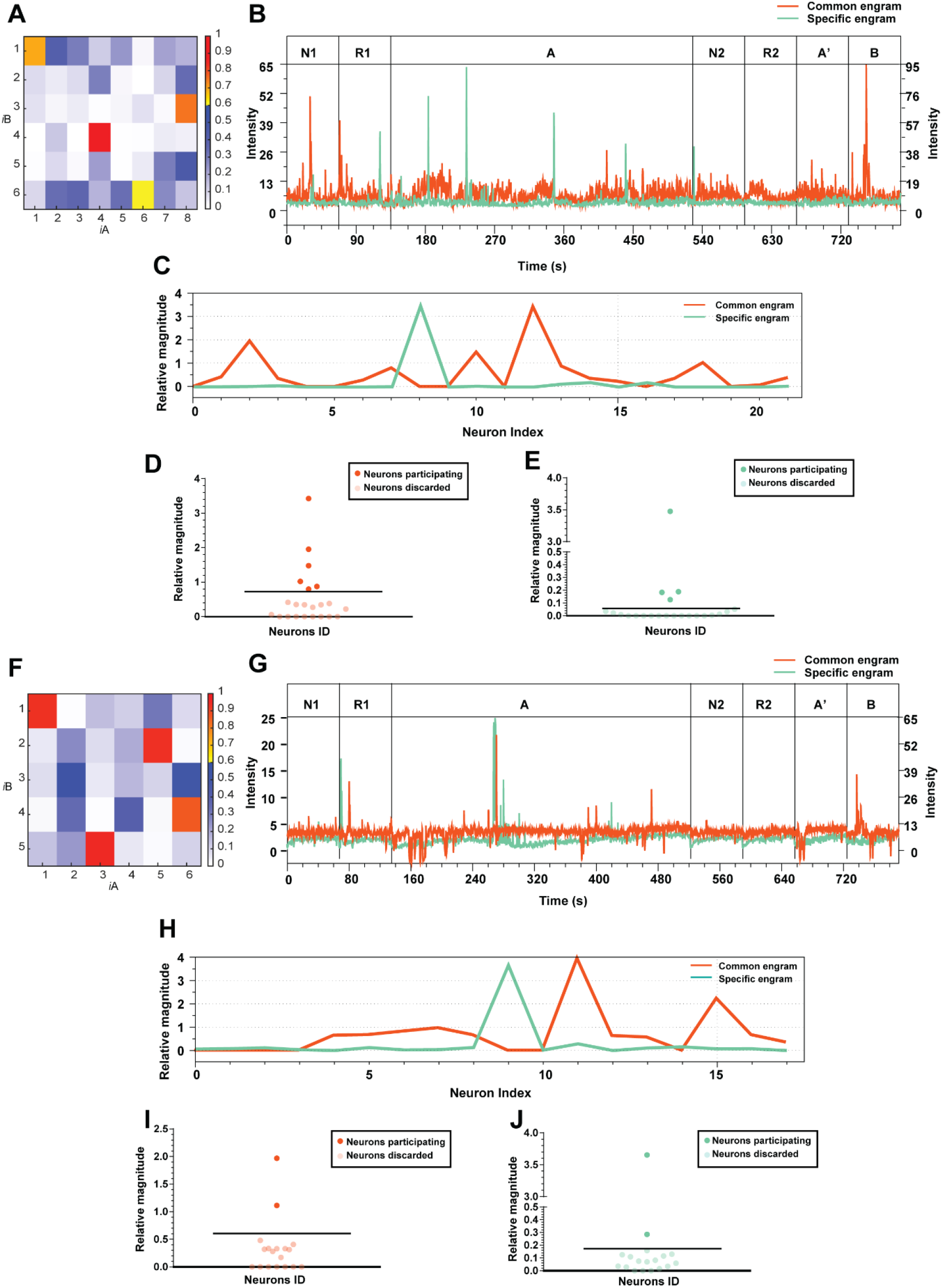
Common and specific engram ensemble activity in all sessions. (A) Dot product analysis (*i*) of NMF-extracted ensembles in engram cells in sessions A and B. (B) Representative ensemble of common engrams (yellow) and specific engrams (light green) across all sessions. The intensity of common engram ensembles is represented on the primary y-axis, and the intensity of specific engram ensembles is represented on the secondary y-axis. (C) Relative magnitude of all neurons participating in common and specific engram ensembles, shown in yellow and light green, respectively. (D & E) Graph showing actively participating neurons in common (yellow) and specific (light green) engram ensembles created by showing those neurons having activity above double the median of the ensemble activity. The line represents double the median. (F-J) Same as (A-D) from a different mouse.

**Suppl. Fig. 10.**
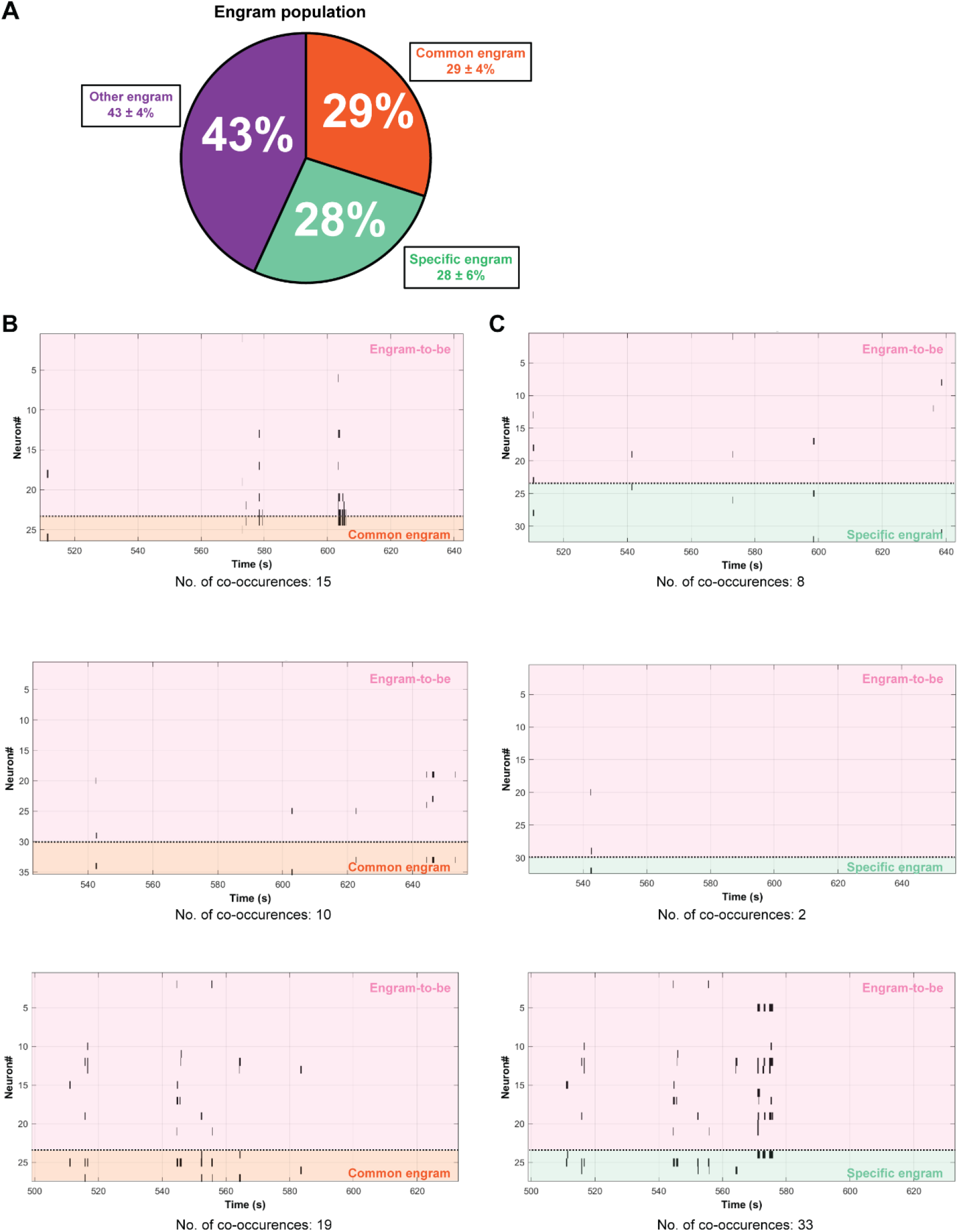
Engram-to-be co-occurrences with common and specific engram populations during postlearning sleep (N2 & R2) sessions. (A) Classification of engram population into common (29%), specific (28%) or other (43%) engram. (B-D) Representative images of number of co-occurrences between engram-to-be with common engram (left) and engram-to-be with specific engram (right) during postlearning sleep sessions (NREM and REM) each tick at certain time point in both 2 populations is counted as a single co-occurrence.

**Suppl. Fig. 11.**
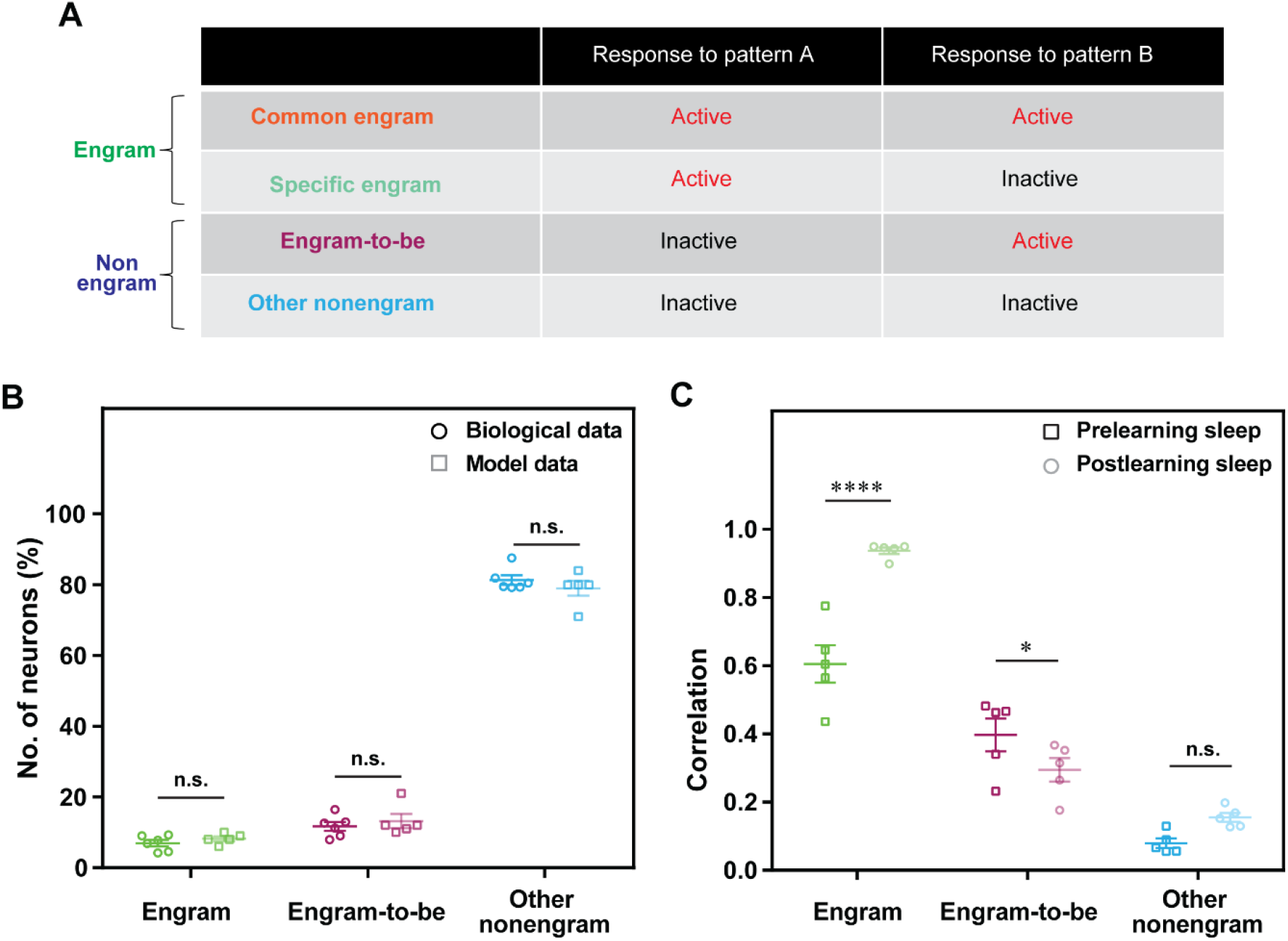
The neural mathematical model follows the biological data. (A) Classification of cell criteria in the neural model data in reference to the biological data. (B) Number of neurons in each cell category in both the biological data (dark colored) and neural model data (light colored). Statistical comparisons were made between the biological data (n = 6) and model data (n = 5) using two-way ANOVA, F (1, 27) = 0.01302, p = 0.9100 (biological data vs model data), Sidak’s multiple comparisons test, p = 0.9047 (Engram), p = 0.8527 (engram-to-be), p = 0.5966 (other nonengram). n.s., nonsignificant. Data represent the mean ± s.e.m. (C) Pearson correlation in the sleep plasticity OFF model in pre- (light colored) and postlearning sleep (dark colored) in engram cells (green), engram-to-be cells (pink), and other nonengrams (light blue). n = 5, statistical comparisons were made between the prelearning sleep and postlearning sleep models using two-way repeated-measures ANOVA, F (1, 4) = 27.99, p = 0.0061 (prelearning sleep vs postlearning sleep), Sidak’s multiple comparisons test, p < 0.0001 (Engram), p = 0.0197 (engram-to-be), p = 0.0794 (other nonengram). ^****^p < 0.0001, ^*^p < 0.05, n.s., nonsignificant. Data represent the mean ± s.e.m.

**Suppl. Fig. 12.**
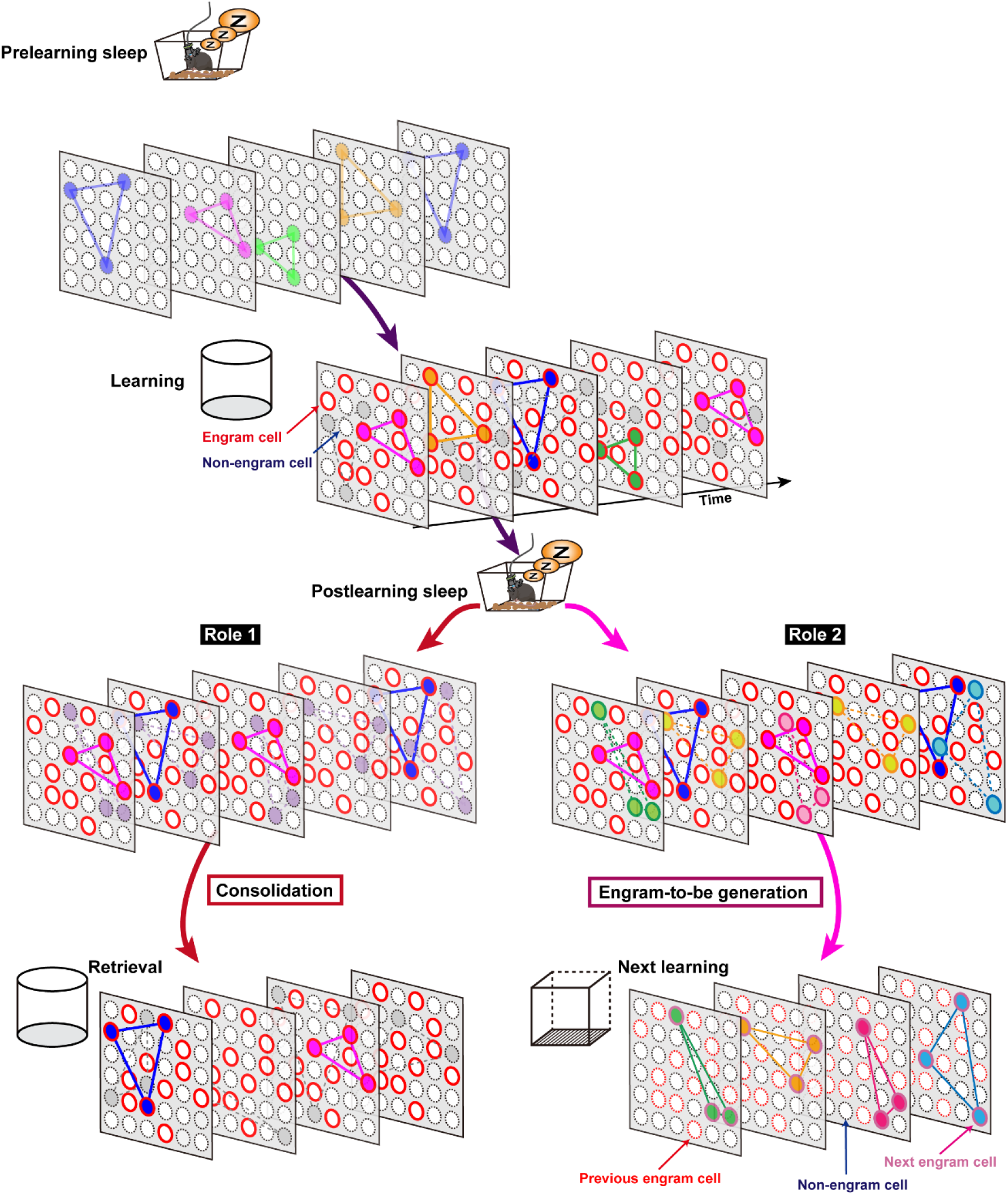
Schematic model showing ensembles activities across memory processing stages. Ensembles organized and activated in a synchronous manner during prelearning sleep preactivated ensembles are later recruited in upcoming engram. Cell population consisting of engram (red circles) and nonengram cells (dotted blue circles). During postlearning sleep, there are two simultaneous processes occurring; engram ensembles activated during learning are then consolidated and later reactivated during retrieval (Role 1). Synaptic plasticity mechanisms, such as synaptic scaling and synaptic depression (LTD) during sharp wave ripples, lead to emergence of new ensembles that are coactivated with previous engram cells as a mechanism for preparing future engram cells for next learning (Role 2). Cell population after a given learning experience consisting of previous engram (dotted red circles), nonengram (dotted blue circles) and next engram cells (pink circles).

**Supplementary Table 1:** Number of engram, engram-to-be and other non-engram cells detected from animals used in this study.

| ID/ Cell type | Engram | Engram-to-be | Other non-engram | Total | Data are used for |
| --- | --- | --- | --- | --- | --- |
| **ID 1** | 19 | 23 | 163 | 205 | Fig. 1E, Fig. 2B, C, Fig. 3C, D, E, Fig.4 C-F, S Fig.1 B-D, S Fig.2 B-G, S Fig. 3C, S Fig.4A, B, S Fig. 5D, E, S Fig.7B, S Fig. 8D, S Fig 10A, B, S Fig 11B, C |
| **ID 2** | 18 | 30 | 183 | 231 | Fig. 1E, Fig. 2B & C, Fig. 3C, D, E, Fig.4A, C-F, S Fig.2F, G, S Fig. 3C, S Fig. 5D, E, S Fig.4A, B, S Fig.7B, S Fig. 8D, S Fig 9F-J, S Fig 10A, C, S Fig 11B, C |
| **ID 3** | 22 | 22 | 199 | 243 | Fig. 1E, Fig. 2B, C, Fig. 3C, D, E, Fig. 4 C-F, S Fig.2 F, G, S Fig. 3C, S Fig.4A, B, S Fig. 5C-E, S Fig.7B, S Fig. 8D, S Fig 9A-E, S Fig 11B, C |
| **ID 4** | 11 | 43 | 207 | 261 | Fig.1E, Fig. 2B, C, Fig. 3C, D, E, S Fig.2 F, G, S Fig. 3C, S Fig.4A, B, S Fig. 5D, E, S Fig. 6E, F, S Fig.7B S Fig. 8D, S Fig 11B, C |
| **ID 5** | 6 | 11 | 70 | 87 | Fig.1E, Fig. 2B, C, Fig. 3C, D, E, S Fig.2 F, G, S Fig. 3C, S Fig.4A, B, S Fig. 5D, E, S Fig.7B, S Fig. 8D, S Fig 11B, C |
| **ID 6** | 13 | 23 | 253 | 289 | Fig.1D, E, Fig. 2B, C, Fig. 3C-F, Fig. 4 C-F, S Fig.2 F, G, S Fig. 3 C-G, S Fig.4A, B, S Fig. 5B, D, E, S Fig.7B, S Fig. 8A-D, S Fig 10A, D |
| **ID 7** | 19 | 70 | 336 | 425 | Fig.3D-F, S Fig.4D, S Fig.6A, B, S Fig.7B |
| **ID 8** | 37 | 314 | 688 | 1039 | Fig.3D-F, Fig.4C-F, S Fig.4D, S Fig.6A, B, S Fig.7B |
| **ID 9** | 8 | N/A | 33 | 41 | S Fig.4D |
| **ID 10** | 7 | 18 | 129 | 154 | Fig.3D-F, S Fig.4D, S Fig.6A, B |
| **Average** | 16 | 56 | 226 | 298 |  |
| **Percentage** | 7 (± 1.5) | 14 (± 2) | 79 (± 2) | 100 | |

## References

1 Silva, A. J., Zhou, Y., Rogerson, T., Shobe, J. & Balaji, J. Molecular and Cellular Approaches to Memory Allocation in Neural Circuits. Science 326, 391–395, doi:doi:10.1126/science.1174519 (2009).

2 Josselyn, S. A. & Tonegawa, S. Memory engrams: Recalling the past and imagining the future. Science 367, eaaw4325, doi:doi:10.1126/science.aaw4325 (2020).

3 Ohkawa, N. et al. Artificial Association of Pre-stored Information to Generate a Qualitatively New Memory. Cell Reports 11, 261–269, doi:10.1016/j.celrep.2015.03.017 (2015).

4 Reijmers, L. G., Perkins, B. L., Matsuo, N. & Mayford, M. Localization of a stable neural correlate of associative memory. Science 317, 1230–1233 (2007).

5 Abdou, K. et al. Synapse-specific representation of the identity of overlapping memory engrams. Science 360, 1227–1231, doi:doi:10.1126/science.aat3810 (2018).

6 Nomoto, M. et al. Cellular tagging as a neural network mechanism for behavioural tagging. Nat Commun 7, 12319, doi:10.1038/ncomms12319 (2016).

7 Goode, T. D., Tanaka, K. Z., Sahay, A. & McHugh, T. J. An Integrated Index: Engrams, Place Cells, and Hippocampal Memory. Neuron 107, 805–820, doi:10.1016/j.neuron.2020.07.011 (2020).

8 Ghandour, K. et al. Orchestrated ensemble activities constitute a hippocampal memory engram. Nature Communications 10, 2637, doi:10.1038/s41467-019-10683-2 (2019).

9 Tanaka, K. Z. et al. The hippocampal engram maps experience but not place. Science 361, 392–397, doi:doi:10.1126/science.aat5397 (2018).

10 Cai, D. J. et al. A shared neural ensemble links distinct contextual memories encoded close in time. Nature 534, 115, doi:10.1038/nature17955 (2016).

11 Rashid, A. J. et al. Competition between engrams influences fear memory formation and recall. Science 353, 383–387, doi:doi:10.1126/science.aaf0594 (2016).

12 Sharpe, M. J., Batchelor, H. M., Mueller, L. E., Gardner, M. P. H. & Schoenbaum, G. Past experience shapes the neural circuits recruited for future learning. Nature Neuroscience 24, 391–400, doi:10.1038/s41593-020-00791-4 (2021).

13 Yiu, A. P. et al. Neurons are recruited to a memory trace based on relative neuronal excitability immediately before training. Neuron 83, 722–735, doi:10.1016/j.neuron.2014.07.017 (2014).

14 Han, J. H. et al. Neuronal competition and selection during memory formation. Science 316, 457–460, doi:10.1126/science.1139438 (2007).

15 Park, S. et al. Neuronal Allocation to a Hippocampal Engram. Neuropsychopharmacology 41, 2987–2993, doi:10.1038/npp.2016.73 (2016).

16 Gouty-Colomer, L. A. et al. Arc expression identifies the lateral amygdala fear memory trace. Molecular Psychiatry 21, 364–375, doi:10.1038/mp.2015.18 (2016).

17 Dragoi, G. & Tonegawa, S. Preplay of future place cell sequences by hippocampal cellular assemblies. Nature 469, 397–401, doi:10.1038/nature09633 (2011).

18 McKenzie, S. et al. Preexisting hippocampal network dynamics constrain optogenetically induced place fields. Neuron 109, 1040-1054.e1047, doi:10.1016/j.neuron.2021.01.011 (2021).

19 Valero, M., Zutshi, I., Yoon, E. & Buzsáki, G. Probing subthreshold dynamics of hippocampal neurons by pulsed optogenetics. Science 375, 570-574, doi:doi:10.1126/science.abm1891 (2022).

20 Aly, M. H., Abdou, K., Okubo-Suzuki, R., Nomoto, M. & Inokuchi, K. Selective engram coreactivation in idling brain inspires implicit learning. Proceedings of the National Academy of Sciences 119, e2201578119. doi:doi:10.1073/pnas.2201578119 (2022).

21 Abdou K N. M., Aly MH, Ibrahim AZ, Choko K, Okubo-Suzuki R, Muramatsu Si, Inokuchi K. Prefrontal coding of learned and inferred knowledge during REM and NREM sleep. Nat. Commun. In press (2024).

22 Huff, M. L., Miller, R. L., Deisseroth, K., Moorman, D. E. & LaLumiere, R. T. Posttraining optogenetic manipulations of basolateral amygdala activity modulate consolidation of inhibitory avoidance memory in rats. Proc Natl Acad Sci U S A 110, 3597–3602, doi:10.1073/pnas.1219593110 (2013).

23 Findlay, G., Tononi, G. & Cirelli, C. The evolving view of replay and its functions in wake and sleep. Sleep advances : a journal of the Sleep Research Society 1, zpab002, doi:10.1093/sleepadvances/zpab002 (2020).

24 Boyce, R., Glasgow, S. D., Williams, S. & Adamantidis, A. Causal evidence for the role of REM sleep theta rhythm in contextual memory consolidation. Science 352, 812-816, doi:doi:10.1126/science.aad5252 (2016).

25 Girardeau, G., Benchenane, K., Wiener, S. I., Buzsáki, G. & Zugaro, M. B. Selective suppression of hippocampal ripples impairs spatial memory. Nature Neuroscience 12, 1222, doi:10.1038/nn.2384 https://www.nature.com/articles/nn.2384#supplementary-information (2009).

26 Norimoto, H. et al. Hippocampal ripples down-regulate synapses. Science 359, 1524-1527, doi:doi:10.1126/science.aao0702 (2018).

27 Wagner, U., Gais, S., Haider, H., Verleger, R. & Born, J. Sleep inspires insight. Nature 427, 352–355, doi:10.1038/nature02223 (2004).

28 Lewis, P. A., Knoblich, G. & Poe, G. How Memory Replay in Sleep Boosts Creative Problem-Solving. Trends in Cognitive Sciences 22, 491–503, doi:10.1016/j.tics.2018.03.009 (2018).

29 Lacaux, C. et al. Sleep onset is a creative sweet spot. Science Advances 7, eabj5866. doi:doi:10.1126/sciadv.abj5866 (2021).

30 Pfeiffer, B. E. & Foster, D. J. Hippocampal place-cell sequences depict future paths to remembered goals. Nature 497, 74–79, doi:10.1038/nature12112 (2013).

31 Wu, C.-T., Haggerty, D., Kemere, C. & Ji, D. Hippocampal awake replay in fear memory retrieval. Nature Neuroscience 20, 571–580, doi:10.1038/nn.4507 (2017).

32 Addis, D. R., Wong, A. T. & Schacter, D. L. Remembering the past and imagining the future: Common and distinct neural substrates during event construction and elaboration. Neuropsychologia 45, 1363–1377, doi:10.1016/j.neuropsychologia.2006.10.016 (2007).

33 Szpunar, K. K., Watson, J. M. & McDermott, K. B. Neural substrates of envisioning the future. Proceedings of the National Academy of Sciences 104, 642, doi:10.1073/pnas.0610082104 (2007).

34 Schacter, D. L., Addis, D. R. & Buckner, R. L. Remembering the past to imagine the future: the prospective brain. Nature Reviews Neuroscience 8, 657–661, doi:10.1038/nrn2213 (2007).

35 Hassabis, D., Kumaran, D., Vann, S. D. & Maguire, E. A. Patients with hippocampal amnesia cannot imagine new experiences. Proceedings of the National Academy of Sciences 104, 1726-1731, doi:doi:10.1073/pnas.0610561104 (2007).

36 Dragoi, G. & Tonegawa, S. Selection of preconfigured cell assemblies for representation of novel spatial experiences. Philosophical transactions of the Royal Society of London. Series B, Biological sciences 369, 20120522, doi:10.1098/rstb.2012.0522 (2014).

37 Asai, H. et al. Pcdhβ deficiency affects hippocampal CA1 ensemble activity and contextual fear discrimination. Molecular Brain 13, 7, doi:10.1186/s13041-020-0547-z (2020).

38 Ghosh, K. K. et al. Miniaturized integration of a fluorescence microscope. Nat Methods 8, 871–878, doi:10.1038/nmeth.1694 (2011).

39 Tsutsui, H., Karasawa, S., Shimizu, H., Nukina, N. & Miyawaki, A. Semi-rational engineering of a coral fluorescent protein into an efficient highlighter. EMBO Reports 6, 233–238, doi:10.1038/sj.embor.7400361 (2005).

40 Takekawa, T. et al. Automatic sorting system for large calcium imaging data. bioRxiv, doi:10.1101/215145 (2017).

41 Burnham, K. P. & Anderson, D. R. Model selection and multimodel inference: a practical information-theoretic approach. (Springer Science & Business Media, 2003).

42 Lee, D. D. & Seung, H. S. in Advances in neural information processing systems. 556–562.

43 Sehgal, M. et al. Memory allocation mechanisms underlie memory linking across time. Neurobiology of Learning and Memory 153, 21–25, doi:10.1016/j.nlm.2018.02.021 (2018).

44 Choi, J.-H. et al. Interregional synaptic maps among engram cells underlie memory formation. Science 360, 430 (2018).

45 Shepherd, J. D. et al. Arc/Arg3.1 Mediates Homeostatic Synaptic Scaling of AMPA Receptors. Neuron 52, 475–484, doi:10.1016/j.neuron.2006.08.034 (2006).

46 Béïque, J.-C., Na, Y., Kuhl, D., Worley, P. F. & Huganir, R. L. Arc-dependent synapse-specific homeostatic plasticity. Proceedings of the National Academy of Sciences 108, 816-821, doi:doi:10.1073/pnas.1017914108 (2011).

47 Okuno, H. et al. Inverse Synaptic Tagging of Inactive Synapses via Dynamic Interaction of Arc/Arg3.1 with CaMKIIβ. Cell 149, 886–898, doi:10.1016/j.cell.2012.02.062 (2012).

48 Grosmark, A. D. & Buzsáki, G. Diversity in neural firing dynamics supports both rigid and learned hippocampal sequences. Science 351, 1440-1443, doi:doi:10.1126/science.aad1935 (2016).

49 Malvache, A., Reichinnek, S., Villette, V., Haimerl, C. & Cossart, R. Awake hippocampal reactivations project onto orthogonal neuronal assemblies. Science 353, 1280–1283, doi:10.1126/science.aaf3319 (2016).

50 Miyawaki, H. & Mizuseki, K. De novo inter-regional coactivations of preconfigured local ensembles support memory. Nature Communications 13, 1272, doi:10.1038/s41467-022-28929-x (2022).

51 Vaz, A. P., Wittig, J. H., Inati, S. K. & Zaghloul, K. A. Backbone spiking sequence as a basis for preplay, replay, and default states in human cortex. Nature Communications 14, doi:10.1038/s41467-023-40440-5 (2023).

52 Holtmaat, A. & Caroni, P. Functional and structural underpinnings of neuronal assembly formation in learning. Nat Neurosci 19, 1553–1562, doi:10.1038/nn.4418 (2016).

53 Huszár, R., Zhang, Y., Blockus, H. & Buzsáki, G. Preconfigured dynamics in the hippocampus are guided by embryonic birthdate and rate of neurogenesis. Nature Neuroscience 25, 1201–1212, doi:10.1038/s41593-022-01138-x (2022).

54 Mocle, A. J. et al. Excitability mediates allocation of pre-configured ensembles to a hippocampal engram supporting contextual conditioned threat in mice. Neuron 112, 1487-1497.e1486, doi:10.1016/j.neuron.2024.02.007 (2024).

55 Buzsáki, G. The Brain from Inside Out. (Oxford University Press, 2019).

56 van de Ven, G. M., Trouche, S., McNamara, C. G., Allen, K. & Dupret, D. Hippocampal Offline Reactivation Consolidates Recently Formed Cell Assembly Patterns during Sharp Wave-Ripples. Neuron 92, 968–974, doi:10.1016/j.neuron.2016.10.020 (2016).

57 Sun, X. et al. Functionally Distinct Neuronal Ensembles within the Memory Engram. Cell 181, 410-423.e417, doi:10.1016/j.cell.2020.02.055 (2020).

58 Nambu, M. F., Lin, Y.-J., Reuschenbach, J. & Tanaka, K. Z. What does engram encode?: Heterogeneous memory engrams for different aspects of experience. Current Opinion in Neurobiology 75, 102568, doi:10.1016/j.conb.2022.102568 (2022).

59 Udakis, M., Pedrosa, V., Chamberlain, S. E. L., Clopath, C. & Mellor, J. R. Interneuron-specific plasticity at parvalbumin and somatostatin inhibitory synapses onto CA1 pyramidal neurons shapes hippocampal output. Nature Communications 11, 4395, doi:10.1038/s41467-020-18074-8 (2020).

60 Geiller, T. et al. Local circuit amplification of spatial selectivity in the hippocampus. Nature 601, 105–109, doi:10.1038/s41586-021-04169-9 (2022).

61 Rolotti, S. V. et al. Local feedback inhibition tightly controls rapid formation of hippocampal place fields. Neuron 110, 783-794.e786, doi:10.1016/j.neuron.2021.12.003 (2022).

62 Sato, M. et al. Distinct Mechanisms of Over-Representation of Landmarks and Rewards in the Hippocampus. Cell Reports 32, 107864, doi:10.1016/j.celrep.2020.107864 (2020).

63 Matsuo, N., Reijmers, L. & Mayford, M. Spine-Type-Specific Recruitment of Newly Synthesized AMPA Receptors with Learning. Science 319, 1104 (2008).

64 Sato, M. et al. Dynamic embedding of salience coding in hippocampal spatial maps. bioRxiv, doi:10.1101/266767 (2018).

65 Barretto, R. P. et al. Time-lapse imaging of disease progression in deep brain areas using fluorescence microendoscopy. Nat Med 17, 223–228, doi:nm.2292[pii]10.1038/nm.2292 (2011).

66 Ziv, Y. et al. Long-term dynamics of CA1 hippocampal place codes. Nat Neurosci 16, 264–266, doi:10.1038/nn.3329 (2013).

67 Jimenez, J. C. et al. Anxiety Cells in a Hippocampal-Hypothalamic Circuit. Neuron 97, 670-683.e676, doi:10.1016/j.neuron.2018.01.016 (2018).

68 Peyrache, A., Benchenane, K., Khamassi, M., Wiener, S. I. & Battaglia, F. P. Principal component analysis of ensemble recordings reveals cell assemblies at high temporal resolution. J Comput Neurosci 29, 309–325, doi:10.1007/s10827-009-0154-6 (2010).

69 Sasaki, T., Kimura, R., Tsukamoto, M., Matsuki, N. & Ikegaya, Y. Integrative spike dynamics of rat CA1 neurons: a multineuronal imaging study. J Physiol 574, 195–208, doi:jphysiol.2006.108480[pii]10.1113/jphysiol.2006.108480 (2006).

70 Carrillo-Reid, L., Miller, J. E., Hamm, J. P., Jackson, J. & Yuste, R. Endogenous sequential cortical activity evoked by visual stimuli. J Neurosci 35, 8813–8828, doi:35/23/8813[pii]10.1523/JNEUROSCI.5214-14.2015 (2015).

71 Middleton, S. J. et al. Altered hippocampal replay is associated with memory impairment in mice heterozygous for the Scn2a gene. Nature Neuroscience 21, 996–1003, doi:10.1038/s41593-018-0163-8 (2018).

72 Trouche, S. et al. Recoding a cocaine-place memory engram to a neutral engram in the hippocampus. Nature Neuroscience 19, 564–567, doi:10.1038/nn.4250 (2016).

73 Mahalanobis, P. On tests and measures of group divergence I. Theoretical formulae. Journal of the Asiatic Society of Bengal 26, 541–588 (1930).

